# Toll-Like-Receptor 5 protects against pulmonary fibrosis by reducing lung dysbiosis

**DOI:** 10.1101/2024.04.30.591719

**Authors:** Yosuke Sakamachi, Emma Wiley, Alma Solis, Collin G Johnson, Xianglin Meng, Salik Hussain, Jay H Lipinski, David N O’Dwyer, Thomas Randall, Jason Malphurs, Brian Papas, Benjamin G Wu, Yonghua Li, Matthias Kugler, Sanya Mehta, Carol S Trempus, Seddon Y Thomas, Jian-Liang Li, Lecong Zhou, Peer W Karmaus, Michael B Fessler, John A McGrath, Kevin Gibson, Daniel J Kass, Anatoli Gleiberman, Avram Walts, Rachele Invernizzi, Phil L Molyneaux, Ivana V Yang, Yingze Zhang, Naftali Kaminski, Leopoldo N Segal, David A Schwartz, Andrei V Gudkov, Stavros Garantziotis

## Abstract

Idiopathic pulmonary fibrosis (IPF) is a devastating pulmonary disease with no curative treatment other than lung transplantation. IPF results from maladaptive responses to lung epithelial injury, but the underlying mechanisms remain unclear. Here, we show that deficiency in the innate immune receptor, toll-like receptor 5 (TLR5), is associated with IPF in humans and with increased susceptibility to epithelial injury and experimental fibrosis in mice, while activation of lung epithelial TLR5 through a synthetic flagellin analogue protects from experimental fibrosis. Mechanistically, epithelial TLR5 activation induces antimicrobial gene expression and ameliorates dysbiosis after lung injury. In contrast, TLR5 deficiency in mice and IPF patients is associated with lung dysbiosis. Elimination of the microbiome in mice through antibiotics abolishes the protective effect of TLR5 and reconstitution of the microbiome rescues the observed phenotype. In aggregate, TLR5 deficiency is associated with IPF and dysbiosis in humans and in the murine model of pulmonary fibrosis. Furthermore, TLR5 protects against pulmonary fibrosis in mice and this protection is mediated by effects on the microbiome.

**One-sentence summary:** Deficiency in the innate immune receptor TLR5 is a risk factor for pulmonary fibrosis, because TLR5 prevents microbial dysbiosis after lung injury.

## INTRODUCTION

Idiopathic pulmonary fibrosis (IPF) is a progressive, fatal lung disease that poses a serious health concern due to its high mortality and paucity of therapeutic options (*1*). Although the pathogenesis remains elusive, it is now widely accepted that IPF is caused by a maladaptive response to lung injury, leading to severe alveolar destruction and irreversible deposition of extracellular matrix in the interstitium (*2*). A growing body of evidence suggests that polymorphisms in genes involved in epithelial integrity and host defense contribute to IPF susceptibility (*3*).

Since host defense has evolved to recognize pathogen-associated molecular patterns, it is reasonable to investigate a connection between IPF and the microbiome. Indeed, microbial dysbiosis was recently found to be associated with IPF progression and mortality (*4–6*), while in mouse models of pulmonary fibrosis, dysbiosis precedes fibrosis and is necessary for its development (*5, 7*). Despite the emerging association between microbial dysbiosis and IPF, genetic factors regulating the microbiome in the context of IPF are yet to be described.

Toll-like receptor 5 (TLR5) is an innate immune receptor recognizing bacterial flagellin (*8*) and is involved in host defense and microbiome modulation (*9*), through which it promotes epithelial integrity. *Tlr5*-deficient mice have increased intestinal injury and develop spontaneous colitis due to dysregulation of the intestinal microbiome (*10*), which mediates a breach of the intestinal mucosal barrier, causing inflammation and tissue damage (*11*). In humans, the minor allele of *TLR5* gene single nucleotide polymorphism rs5744168 (minor allele frequency 5.6%) introduces a STOP codon after the transmembrane domain of *TLR5*, thus in effect creating a decoy receptor and disrupting TLR5 signaling (*12, 13*). Carriers of this polymorphism have increased susceptibility to infections of the airway (*12*) and urinary tract (*14*). Collectively, these findings underscore the central role of TLR5 in the immune response to microbiota.

Because TLR5 activation regulates epithelial integrity after injury (*15, 16*) and regulates microbiome abundance across epithelial surfaces, and since other host defense genes have been implicated in genetic susceptibility to IPF, we hypothesized that TLR5 may play a role in lung fibrosis. Here, we first explored the association of the functionally deficient minor allele of rs5744168 with IPF in human patients and investigated TLR5 expression in healthy and fibrotic human lungs. We then examined the role of genetic *Tlr5* deficiency in the susceptibility to lung fibrosis and whether pharmacological activation of TLR5 ameliorates fibrosis. Finally, we determined the interactions of TLR5 with the lung microbiome in lung fibrosis and explored associations of the minor allele of rs5744168 with dysbiosis in lungs of IPF patients.

## RESULTS

### Case-control genotypes in the cohorts

Since rs5744168 is not included in any of the major GWAS panels, we genotyped our cohorts for this SNP *de novo*. Our discovery cohort included 277 IPF patients and 397 lung-healthy controls from the University of Pittsburgh and our validation cohort was a nation-wide sample of 833 IPF cases collected at the University of Colorado and 2505 lung-healthy controls derived from the COPDGene cohort (**Supplementary Table 1**). We tested our target TLR5 SNP rs5744168 and the well-described MUC5B SNP rs35705059 as a positive reference control. Numbers varied slightly by SNP within each dataset, depending on genotyping success. In the discovery dataset, rs5744168 did not meet the criterion for Hardy-Weinberg equilibrium (HWE) in unaffected control subjects (HWE, p = 5.143E-07); it met HWE in IPF cases (p = 0.2348), while rs35705950 met HWE in cases and controls (p = 1.0). The discovery cohort HWE results probably reflect the relatively small sample size. In the validation dataset both rs5744168 and rs35705950 met HWE (p = 0.8523 and 0.003 respectively). In the combined dataset both SNPs met HWE: for rs5744168 p=0.02538, and for rs35705950 p=0.009521. **Supplementary Table 2** provides the genotype breakdown by sample source and SNP, and **Supplementary Table 3** provides the genotype breakdown after combining samples, for subjects with complete genotypes for either rs5744168 or rs35705950: N = 4020 (3858 subjects with rs35705950 and 4012 subjects with rs5744168).

### Established association of IPF with MUC5B rs35705950

We confirmed that our discovery and validation samples are representative of published data, finding a significant additive association of rs35705059 with IPF in both datasets, and in the meta-analysis, with Odds Ratios (ORs) of 5.11 [3.69 – 7.08, p < 0.0001], 8.75 [7.3 – 10.48, p < 0.0001], and 7.70 [6.58 – 9.03, p < 0.0001] for discovery, validation, and meta-analysis respectively (**Table 1**). Comparing cohorts, the METAL heterogeneity test p-value for rs35705950 was significant (p = 0.005). However, the rs35705950 ORs for both the discovery and validation datasets are well within the established range for this SNP (*17*).

**Table 1.** Association of TLR5 (rs5744188: additive and dominant) and MUC5B (rs35705950: additive) SNPs with IPF in the discovery, validation, and meta-analysis cohorts.

| **Cohort** | **rs5744168: additive** |  | **rs35705950: additive** |  |
| --- | --- | --- | --- | --- |
|  | **OR (95% CI)** | **p-value** | **OR (95% CI)** | **p-value** |
| Discovery | 1.20 (0.81-1.77) | 0.36 | 5.11 (3.69 - 7.08) | **<0.0001** |
| Validation | 1.31 (1.06-1.63) | **0.01** | 8.75 (7.3 - 10.48) | **<0.0001** |
| Meta-analysis | 1.28 (1.07-1.55) | **0.009** | 7.70 (6.58-9.03) | **<0.0001** |
| **rs5744168: dominant** |  |  |  |  |
| Discovery | 1.60 (1.01 – 2.51) | **0.04** |  |  |
| Validation | 1.21 (0.96 – 1.54) | 0.10 |  |  |
| Meta-analysis | 1.29 (1.04 – 1.59) | **0.018** |  |  |

### TLR5 rs5744168 is associated with IPF in humans

The ORs and 95% confidence intervals (CI) for the associations of rs35705950 (additive) and rs5744168 (additive and dominant) with IPF are given in **Table 1** for the two cohorts and for the meta-analysis. In the discovery cohort, there was a significant dominant association (OR = 1.60 [1.01 - 2.51], p = 0.04) of rs5744168 with IPF; however, we did not find a significant additive association (OR = 1.20 [95% CI: 0.81 - 1.77], p = 0.36). Of note, there were no homozygote minor allele carriers in the IPF group of this cohort, which may have affected the power of the additive model. In the validation cohort we found a significant additive association of the minor allele of rs5744168 with IPF (OR = 1.31 [1.06 - 1.63], p = 0.01), and a trend for association in the dominant model (OR = 1.21 [0.96 – 1.54], p = 0.10). In the meta-analysis incorporating the discovery and validation datasets, we found a significant additive association of rs5744168 with IPF (OR = 1.28 [1.07 - 1.55], p = 0.009) and a similarly significant dominant association (OR = 1.29 [1.04 – 1.59], p=0.01). The METAL heterogeneity test p-values for both the additive and dominant effects of rs5744168 were non-significant (p = 0.691 and 0.295 respectively). Supplementary Table 2 provides the genotype details for the discovery and validation datasets.

### Interaction and stratification analysis with *MUC5B* rs35705950

Because *TLR5* and *MUC5B* are both host defense genes, we next assessed a possible interaction between the two. Given that rs35705950 has an established additive effect while rs5744168 could be additive or dominant, we examined both the interaction of the additive-coded SNPs, and the interaction of additive rs35705950 with dominant rs5744168, among subjects having both SNPs (N=3850, Supplementary Table 5).

#### Interaction between additive rs35705950 and additive rs5744168

The interaction of the additive-coded SNPs was non-significant (*p* = 0.3599) over all patients, with significant effects for rs35705950 alone (p<0.0001) and rs5744168 alone (p = 0.0008). However, looking at the interaction in subgroups, the additive OR for rs5744168 in the homozygous major stratum of rs35705059 was significant (OR = 1.59 [1.21 - 2.07]) but became increasingly non-significant in the heterozygous and homozygous-rare strata of rs35705059 (OR = 1.30 [0.92 - 1.83] and OR = 1.07 [0.52 – 2.20], respectively), suggesting that the genetic effect of MUC5B dominates over the TLR5 effect. There was a less pronounced effect in MUC5B ORs, depending on TLR5 genotype: OR for rs35705059 were 8.20 [6.93 - 9.71], 6.73 [4.52 - 10.02] and 5.52 [2.46 - 12.35] for the homozygous-major, heterozygous, and homozygous-rare strata of rs5744168, respectively (**Supplementary Table 6**).

#### Interaction of additive rs35705950 and dominant rs5744168

This interaction was also non-significant (p = 0.2473), with significant effects for additive rs35705950 as above (p<0.0001) and for dominant rs5744168 (p = 0.0011). The same diminution of the rs5744168 effect (now dominant) across the strata of additive rs35705950 is seen as with the above model interacting the two additive-coded SNPs (see Supplementary Table 7). Collectively, these results suggest that the *TLR5* polymorphism, rs5744168, is associated with susceptibility to pulmonary fibrosis, and the effect is most significant in patients without additional host-defense variants.

### *TLR5* expression in healthy and IPF lung is found on epithelia and immune cells

We evaluated the expression of *TLR5* in human lungs using single cell RNA sequencing data from healthy, IPF, COPD and other ILD patient lungs, and confirmed TLR5 protein expression via immunohistochemistry. We found that *TLR5* expression was most prominent in epithelial (bronchial and alveolar type 2) and immune cells regardless of disease status (**Fig. 1A, Supplementary Fig. S1**). In IPF lungs, there was immunohistochemical evidence of TLR5 expression in aberrant bronchiolarized alveolar epithelia (**Fig. 1B, C, Supplementary Fig. S2, S3**). Comparing global *TLR5* expression in IPF and control lungs using the LGRC database (*18*), we found that *TLR5* gene expression was overall moderately increased in IPF lungs (**Fig. 1D**).

**Figure 1.**
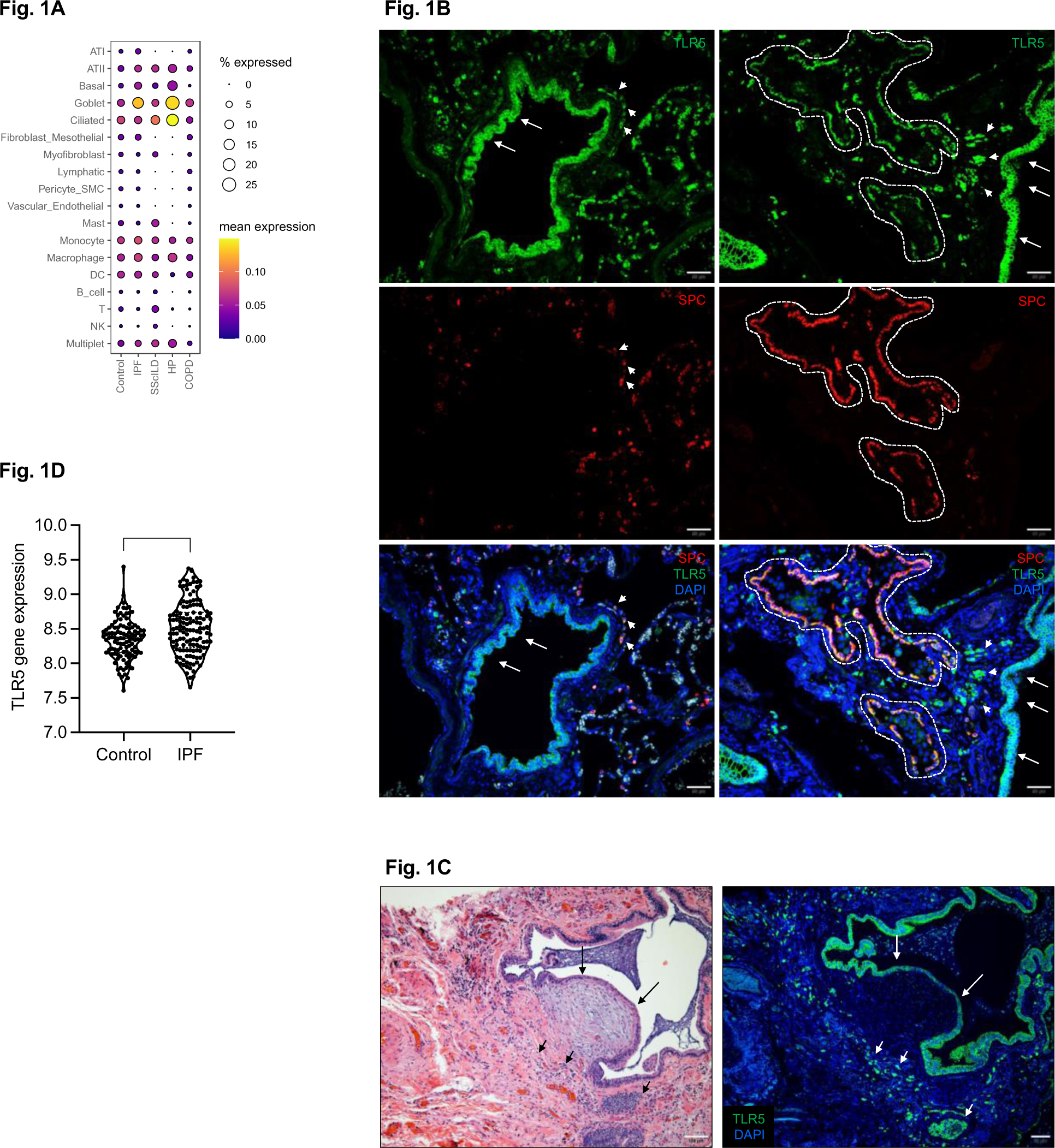
TLR5 SNP expression in healthy and IPF lungs. **(A)** Dot plot derived from scRNA seq data, summarizing *TLR5* gene expression within each major annotated cell type, stratified by disease status. **(B)** Immunohistochemical staining of healthy (left) and IPF lung sections (right); green: TLR5, red: SPC, blue: DAPI. In healthy lungs, note TLR5 staining in bronchial epithelia (arrows) and SPC-positive alveolar Type 2 epithelia (arrowheads). Red blood cells stain false-positive across all stains and are clearly visible as grey/white structures in the merged image. In IPF lungs, note TLR5 staining in aberrant, bronchiolarized SPC-positive alveolar epithelia (discontinuous line surrounding the airway). Representative images shown. **(C)** Left: Hematoxylin and Eosin staining of fibroblastic focus (FF) in IPF lung. Right: immunohistochemical staining of adjacent slide of the same region for TLR5 (green) and DAPI (blue). Note TLR5 staining in aberrant alveolar epithelia over the FF (arrows) and immune cells surrounding the FF (arrowheads). **(D)** Violin plot of TLR5 expression from 108 control and 134 IPF lungs from the Lung Genomics Research Consortium (LGRC). *** p<0.001.

### *Tlr5*-deficient mice are susceptible to bleomycin-induced lung injury and fibrosis

To determine the role of Tlr5 in lung fibrosis, we administered bleomycin to *Tlr5*-wildtype (*Tlr5^+/+^*) and *Tlr5*-deficient (*Tlr5^-/-^*) mice. At 21 days post bleomycin administration, both *Tlr5^-/-^* and *Tlr5^+/+^* mice had significantly higher collagen content in the lung (quantified by hydroxyproline levels) compared to their saline-treated counterparts (**Fig. 2A**). Importantly, *Tlr5^-/-^* mice had significantly higher hydroxyproline levels compared to *Tlr5^+/+^*mice after bleomycin administration (**Fig. 2A**), supported by increased histological evidence of lung fibrosis (**Fig. 2B**) and by increased fibroblast activation (**Fig. 2C**). Over the course of 21 days post bleomycin administration, *Tlr5^-/-^* mice lost significantly more weight (**Fig. 2D**) and had decreased survival (**Fig. 2E**) compared to their *Tlr5^+/+^* littermates. Furthermore, epithelial injury (assayed by protein levels in lung lavage at Day 5 post bleomycin) was significantly increased in *Tlr5^-/-^* mice compared to *Tlr5^+/+^* mice (**Fig. 2F**). Collectively, our results suggest that *Tlr5* protects against lung injury and pulmonary fibrosis in mouse.

**Figure 2.**
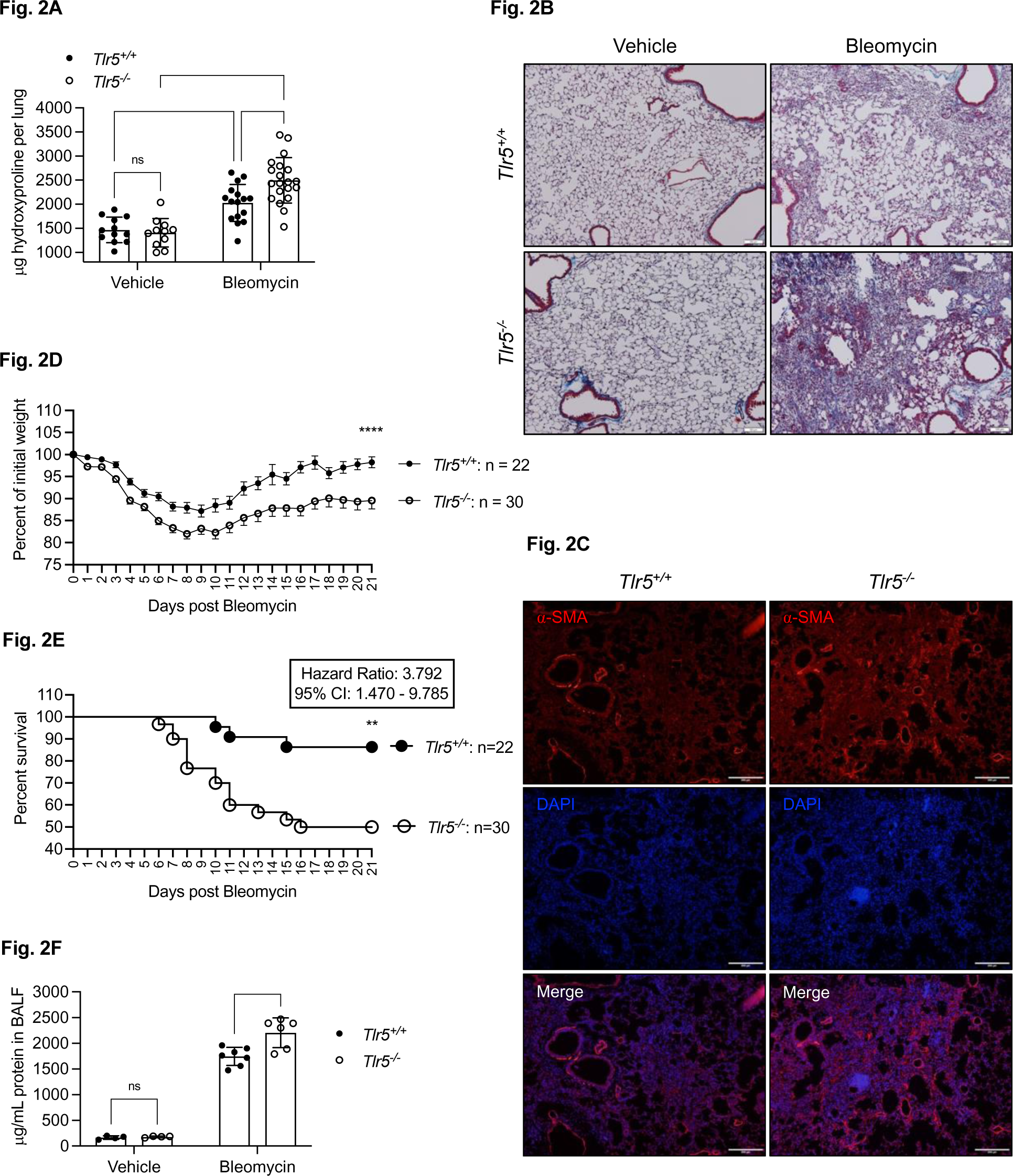
*Tlr5* deficiency exacerbates bleomycin-induced lung injury and fibrosis. **(A).** Quantification of total collagen per lung determined by hydroxyproline assay, 21 days post bleomycin administration in *Tlr5^+/+^* (n=16) and *Tlr5^-/-^* (n=20) mice. Two experiments combined; experiment repeated over 6 times. **(B)** Masson’s Trichrome Staining of *Tlr5^+/+^* and *Tlr5^-/-^* lungs, 21 days post bleomycin treatment. Representative images shown. **(C)** Immunohistochemical staining of *Tlr5^+/+^* and *Tlr5^-/-^* lungs, 21 days post bleomycin administration; red: alpha-smooth muscle actin, blue: DAPI, white: merge. Representative images shown. **(D)** Percent of initial weight of *Tlr5^+/+^* (n=22) and *Tlr5^-/-^* (n=30) mice over 21 days post bleomycin administration. Two experiments combined; experiment repeated over 6 times. **(E)** Percent survival of *Tlr5^+/+^* (n=22) and *Tlr5^-/-^* (n=30) mice over 21 days post bleomycin administration. Two experiments combined; experiment repeated over 6 times. **(F)** Quantification of BALF total proteins in *Tlr5^+/+^* (n=7) and *Tlr5^-/-^* (n=6) mice determined by BCA assay, 5 days post bleomycin administration. Experiment repeated twice. *p<0.05, ** p<0.01, *** p<0.001, **** p<0.0001, one- or two-way ANOVA with post-hoc Holm-Sidak correction or Mantel-Cox Log-rank test as indicated.

### TLR5-agonist ameliorates bleomycin-induced fibrosis through the canonical signaling pathway

Given that *Tlr5^-/-^* mice are susceptible to bleomycin-induced pulmonary fibrosis, we examined whether specific TLR5 activation protects mice against bleomycin-induced fibrosis. To activate TLR5, we dosed mice with a flagellin-derived TLR5-agonist prior to bleomycin administration. Over the course of 21 days post bleomycin administration, wildtype animals dosed with TLR5-agonist lost significantly less weight (**Fig. 3A**) and displayed improved survival (**Fig. 3B**) compared to animals receiving bleomycin and vehicle (saline). Importantly, Tlr5 activation significantly decreased bleomycin-induced fibrosis, as determined by hydroxyproline levels (**Fig. 3C**) and histological evidence (**Fig. 3D**). In further support, lung fibroblast activation was reduced in animals treated with TLR5-agonist/bleomycin compared to animals treated with bleomycin (**Fig. 3E**). TLR5-agonist did not protect *Tlr5^-/-^* mice against bleomycin-induced weight loss or fibrosis, confirming that the observed mechanism is indeed Tlr5-dependent (**Supplementary Fig. 4A, B**). Canonical TLR pathways are mediated via adaptor molecules MYD88 and TRIF. Specifically, TLR5-signaling is mediated by the MYD88 pathway. We found that TLR5-agonist protected *Trif*-deficient mice (*Trif^-/-^*, **Supplementary Fig. 4C, D**) but failed to protect *Myd88*-deficient mice (*Myd88^-/-^*, **Supplementary Fig. 4E, F**) against bleomycin-induced weight loss and fibrosis. Canonical TLR5-MYD88 signaling leads to the activation of NFκB; we found the TLR5-agonist to induce nuclear translocation of the p65 NFκB subunit in lung epithelial cells (**Supplementary Fig. 4G**), 30-minutes to 2-hours post treatment. Collectively, these results suggest that the activation of the canonical TLR5-MYD88 pathway protects against pulmonary fibrosis in mice.

**Figure 3.**
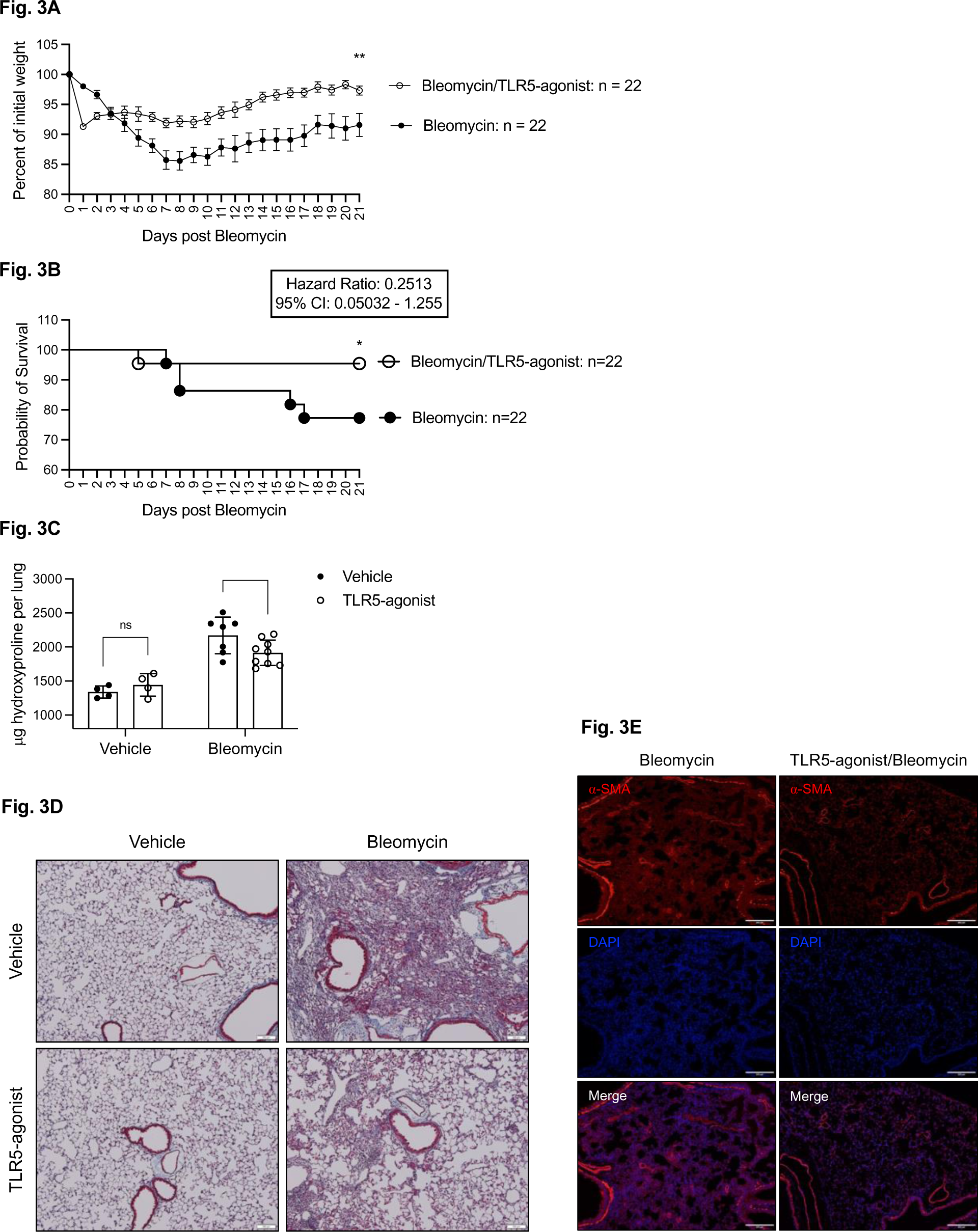
TLR5-agonist protects against bleomycin-induced lung injury and fibrosis. **(A)** Percent of initial weight of vehicle or TLR5-agonist treated animals over 21 days post bleomycin administration (n=22 per group). Experiment repeated over 5 times. **(B)** Percent survival of vehicle and TLR5-agonist mice over 21 days post bleomycin administration (n=22 per group). Experiment repeated over 5 times. **(C)** Quantification of total collagen per lung determined by hydroxyproline assay, 21 days post bleomycin (3U/Kg) administration in vehicle (n=7) and TLR5-agonist treated (n=9) mice. Experiment repeated over 5 times. **(D)** Masson’s Trichrome Staining of vehicle and TLR5-agonist treated lungs, 21 days post bleomycin administration. Representative images shown. **(E)** Immunohistochemical staining of vehicle and TLR5-agonist treated lungs, 21 days post bleomycin administration; red: alpha-smooth muscle actin, blue: DAPI, white: merge. Representative images shown. *p<0.05, one- or two-way ANOVA with post-hoc Holm-Sidak correction or Mantel-Cox Log-rank test as indicated.

We next determined the therapeutic potential of TLR5 activation by administering TLR5-agonist at various time points following bleomycin-injury. TLR5-agonist administration at days 7, 11, 14, and 18 post bleomycin injury significantly reduced lung fibrosis at day 21 with efficacies comparable to Pirfenidone (**Supplementary Fig. 4H**).

### Epithelial *Tlr5* is necessary for the TLR5-agonist protective effect in mouse

We then evaluated which tell type mediates TLR5-protection against bleomycin-induced fibrosis. Because of our finding that TLR5 is primarily expressed in the lung epithelium and in myeloid cells (**Fig. 1A, B**), we generated conditional *Tlr5* knockout mice in the lung epithelium (*Tlr5^Fx/Fx^ Spc-Cre*) or in the myeloid lineage (*Tlr5^Fx/Fx^ LysM-Cre*) and administered bleomycin. We found that *Tlr5* deletion in the myeloid lineage did not result in increased susceptibility to bleomycin-induced fibrosis while *Tlr5* deletion in the lung epithelium resulted in higher collagen content in the lung compared to *Tlr5^+/+^* mice (**Fig. 4A, B**). We next evaluated which cell type mediates the TLR5-agonist protection against bleomycin-induced fibrosis by dosing *Tlr5^Fx/Fx^ Spc-Cre* and *Tlr5^Fx/Fx^ LysM-Cre* mice with TLR5-agonist prior to bleomycin administration. *Tlr5^Fx/Fx^ LysM-Cre* mice had significantly less weight loss and lower levels of hydroxyproline compared to their vehicle-treated *Cre^-^* (*Tlr5^Fx/Fx^*) litter mates, a TLR5-agonist response comparable to that of wild-type mice receiving TLR5 agonist prior to bleomycin. However, *Tlr5^Fx/Fx^ Spc-Cre* failed to respond to the TLR5-agonist (**Fig. 4C, D**), similar to *Tlr5^-/-^* mice. Collectively, these results suggest that epithelial *Tlr5* is important for the observed effects of TLR5-mediated protection against bleomycin-induced pulmonary fibrosis.

**Figure 4.**
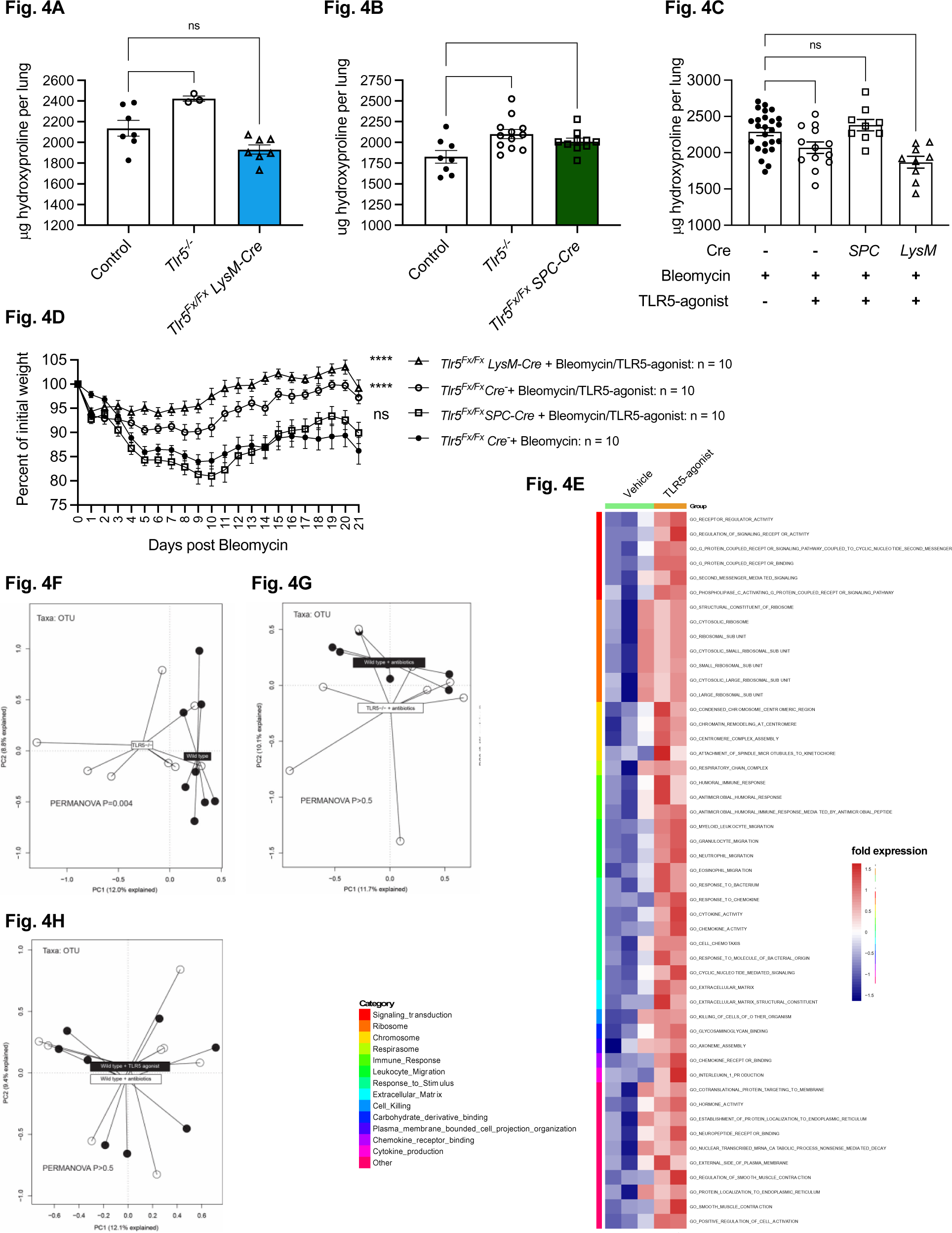
TLR5-agonist acts through lung epithelial *Tlr5* and upregulates antimicrobial genes to prevent dysbiosis. **(A)** Quantification of total collagen per lung, 21 days post bleomycin (3U/Kg) administration, determined by hydroxyproline assay in control (*Tlr5^+/+^*or *Tlr5^Fx/Fx^ Cre^-^*), *Tlr5^-/-^*, and *Tlr5^Fx/Fx^ LysM-Cre* mice. Experiment repeated over 5 times. **(B)** Quantification of total collagen per lung, 21 days post bleomycin (3U/Kg) administration, determined by hydroxyproline assay in control (*Tlr5^+/+^*, *Tlr5^-/-^*), *Tlr5^-/-^*, and *Tlr5^Fx/Fx^ SPC-Cre* mice. Experiment repeated over 5 times. **(C)** Quantification of total collagen per lung determined by hydroxyproline assay in vehicle (n=13) or TLR5-agonist treated mice, 21 days post bleomycin administration (3U/Kg), in *Tlr5^Fx/^*^Fx^ *Cre^-^* (n=13), *Tlr5^Fx/Fx^ Spc*-Cre (n=9), and *Tlr5^Fx/Fx^ LysM-Cre* (n=9) mice. Experiment repeated over 3 times. (A-C) analysis was performed with ANOVA with Holm-Sidak correction, *p<0.05, ** p<0.01, *** p<0.001, **** p<0.0001. **(D)** Percent of initial weight in vehicle (n=9) or TLR5-agonist treated mice over 21 days post bleomycin administration in *Tlr5^Fx/^*^Fx^ *Cre^-^* (n=9), *Tlr5^Fx/Fx^ Spc-Cre* (n=10), and *Tlr5^Fx/Fx^ LysM-Cre* mice (n=10). Experiment repeated over 3 times. **(E)** GSEA analysis of RNA-seq gene signals between vehicle and TLR5-agonist treated groups, using Gene Ontology (GO)-defined gene sets. The Gene sets are grouped according to biological categories; the top 100 GO-defined gene sets that were significantly enriched in the TLR5-agonist group are shown. Each row represents the mean expression of leading genes within the gene set. Complete composition of the groups is listed in Supplementary Table S6 and S7. **(F)** Principal component analysis (PCA) visualizing the altered operational taxonomic units (OTU) in *Tlr5^+/+^* and *Tlr5^-/-^* mice, 7 days post bleomycin administration. **(G)** PCA visualizing the OTU in antibiotic-treated *Tlr5^+/+^* and *Tlr5^-/-^* mice, 7 days post bleomycin administration. **(H)** PCA visualizing the OTU in antibiotic- or TLR5-agonist-treated mice, 7 days post bleomycin administration. (F-H) analysis was performed using permutational multivariate ANOVA.

### Tlr5 modulates the mouse lung microbiome

We then evaluated which pathways are induced after Tlr5 activation by examining gene expression in flow-sorted lung epithelia 36-hours after bleomycin exposure, comparing treatment with the TLR5-agonist or vehicle. We found that most of the differentially expressed pathways in TLR5-agonist-treated mouse lungs were associated with antimicrobial activity and host defense (**Fig. 4E, Supplementary Tables 8, 9**). Given that *Tlr5^Fx/Fx^ Spc-Cre* mice were susceptible to bleomycin-induced fibrosis and did not respond to TLR5-agonism, we examined the gene expression in flow-sorted *Tlr5^Fx/Fx^ Spc-Cre* lung epithelia after TLR5-agonist or vehicle treatment. Indeed, pathways associated with antimicrobial activity and host defense were blunted in *Tlr5*-deficient lung epithelia (**Supplementary Table 10**).

These results suggested that the antifibrotic *Tlr5* activity in the lung may involve modulation of the lung microbiome. We therefore evaluated the lung microbiome in association with TLR5 activity. We found that *Tlr5^-/-^* mouse lungs had significantly dysbiotic microbiome compared to *Tlr5^+/+^* lungs after bleomycin exposure (**Fig. 4F**), which was ameliorated by antibiotic treatment (vancomycin, neomycin, metronidazole, and ampicillin as described in the methods section; **Fig. 4F**). Interestingly, treatment with the TLR5-agonist had similar effects on the murine lung microbiome after bleomycin exposure as treatment with antibiotics (**Fig. 4G**).

To determine whether the lung microbiome can activate lung TLR5, we first evaluated lung *Tlr5* expression in response to flagellin. Compared to naïve specific-pathogen-free (SPF) mouse lungs, lung *Tlr5* expression was decreased after daily dosing with flagellin for 3 days (**Supplementary Fig. 5A**), presumably as a negative feedback mechanism. If negative feedback mechanisms decrease Tlr5 expression after flagellin exposure, we reasoned that elimination of the microbiome in germ-free mice will upregulate *Tlr5* expression. We compared *Tlr5* expression in germ-free (GF) and SPF mice purchased from the same vendor and found that the effect of the microbiome on lung *Tlr5* expression (SPF mice compared to GF mice) was similar to the effect seen after flagellin exposure (**Supplementary Fig. 5B**), suggesting that the native microbiome activates Tlr5 in the murine lung.

Tlr5 protection against fibrosis is mediated by amelioration of dysbiosis after bleomycin exposure.

To evaluate whether Tlr5 protection against fibrosis involves the microbiome, we treated *Tlr5^-/-^* and *Tlr5^+/+^* mice with a quadruple antibiotic cocktail (ABX) of Ampicillin, Vancomycin, Neomycin, and Metronidazole for 4-weeks prior to bleomycin-administration. We found that *Tlr5* genotype did not impact the degree of fibrosis in antibiotic-treated mice, in contrast to non-antibiotic-treated mice (**Fig. 5A**). When we reconstituted the microbiome after antibiotic treatment, we were able to restore the *Tlr5* protective phenotype after bleomycin administration (**Fig. 5B, Supplementary Fig. 5A**). Importantly, treatment with TLR5-agonist conferred no additional benefit over the treatment with antibiotics in mice exposed to bleomycin, suggesting that the protective effect of TLR5-agonism requires microbiota (**Fig. 5C**). It has been reported that dysbiosis precedes lung injury and persists in the fibrotic lung in the bleomycin-induced fibrosis model (*5*), suggesting that antibiotic-treatment alone may be sufficient in reducing the severity of response to bleomycin in ABX/TLR5-agonist treated mice. To determine whether the severity of response to bleomycin contributed to the observed effect, we administered a high dose of bleomycin (4U/Kg) in combination with ABX and/or TLR5-agonist and compared lung fibrosis. We found that both antibiotic-treatment and TLR5-agonism reduced lung fibrosis in animals treated with the high dose of bleomycin (**Fig. 5D**). Importantly, TLR5-agonism conferred no additional benefit in antibiotic-treated animals exposed to high dose of bleomycin, suggesting that the protective effect of TLR5-agonism indeed requires microbiota (**Fig. 5D**). In aggregate, our data support that the *a priori* presence of microbiome is necessary for *Tlr5*-mediated protection against fibrosis.

**Figure 5.**
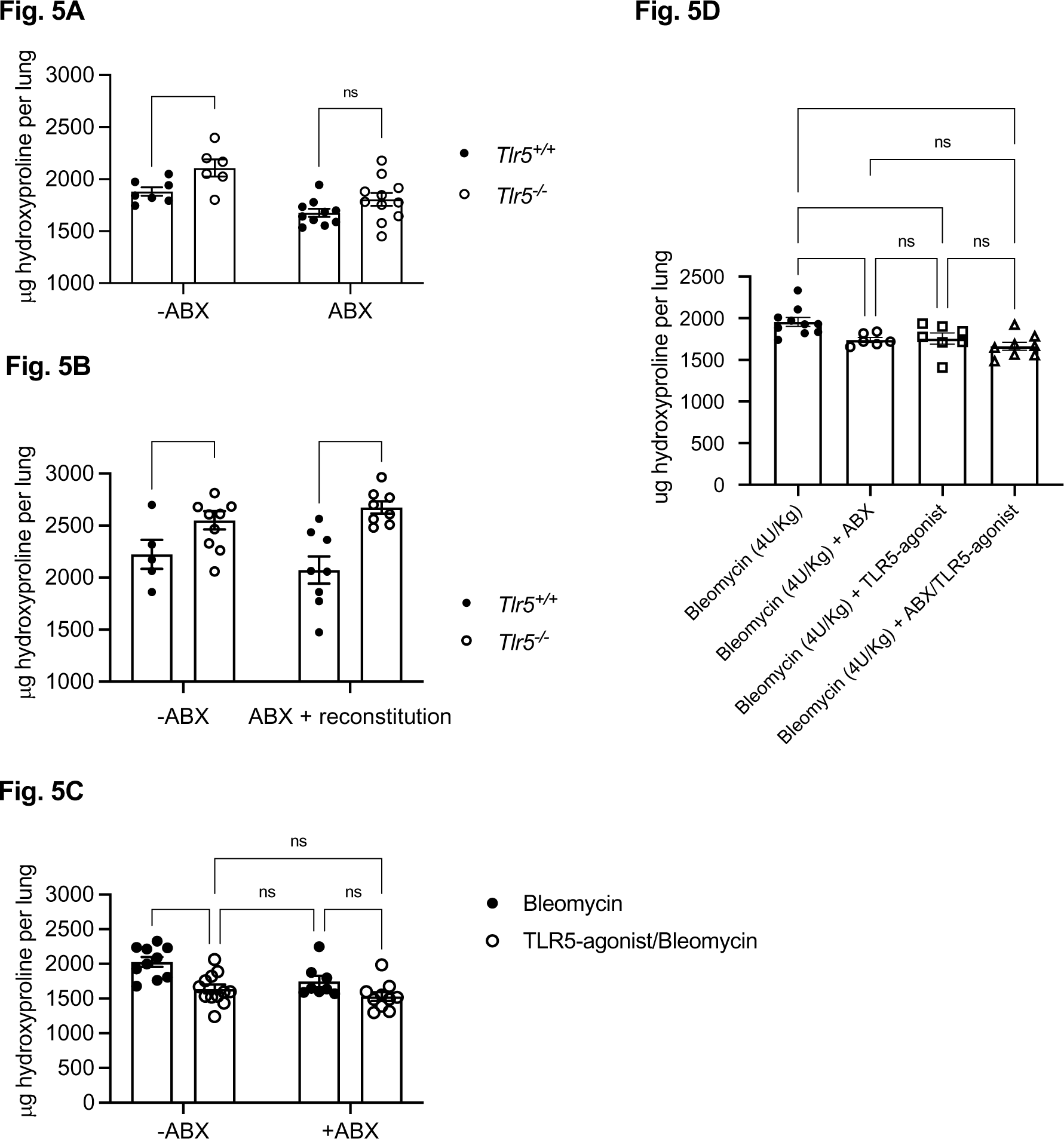
TLR5 activation modifies lung microbiome and TLR5 effect on fibrosis depends on the presence of microbiome. **(A)** Quantification of total collagen per lung determined by hydroxyproline assay in vehicle or antibiotic-treated *Tlr5^+/+^* and *Tlr5^-/-^* mice, 14 days post bleomycin (3U/Kg) administration. Experiment repeated over 3 times. **(B)** Quantification of total collagen per lung determined by hydroxyproline assay in vehicle or antibiotic-reconstituted *Tlr5^+/+^* and *Tlr5^-/-^* mice, 21 days post bleomycin administration (3U/Kg). **(C)** Quantification of total collagen per lung determined by hydroxyproline assay in vehicle or antibiotics, vehicle or TLR5-agonist treated mice, 21 days post bleomycin administration (3U/Kg). (D) Quantification of total collagen per lung determined by hydroxyproline assay in animals treated with high dose (4U/Kg) of bleomycin, 21 days post administration. Analysis was performed with ANOVA with Holm-Sidak correction, *p<0.05, ** p<0.01, *** p<0.001.

### TLR5 deficiency is associated with altered lung lavage microbiome profile in human IPF patients

We last questioned whether TLR5 influences the lung microbiome of IPF patients. To address this, we examined the BALF microbiome of rs5744168 major allele homozygote (N = 85) and minor-allele carrier (N = 18) IPF patients. Bacterial burden was comparable between our non-carrier and carrier cohorts (**Supplementary Fig. 7A**). However, we found a significant reduction in microbial diversity in rs5744168 carriers compared to major allele homozygotes (**Fig. 6A**). At the phylum level, Proteobacteria were increased, while Bacteroidetes were decreased in rs5744168 minor allele carriers (**Fig. 6B**). At the genus level we observed an increase in *Haemophilus, Staphylococcus,* and *Neisseria*, which are associated with IPF disease profession, with a concomitant decrease in *Prevotella* (and trend for *Veillonella* in adjusted analysis) in rs5744168 minor allele carriers (**Fig. 6C**). Collectively, our results suggest that TLR5 deficiency is associated with increased dysbiosis among IPF patients.

**Figure 6.**
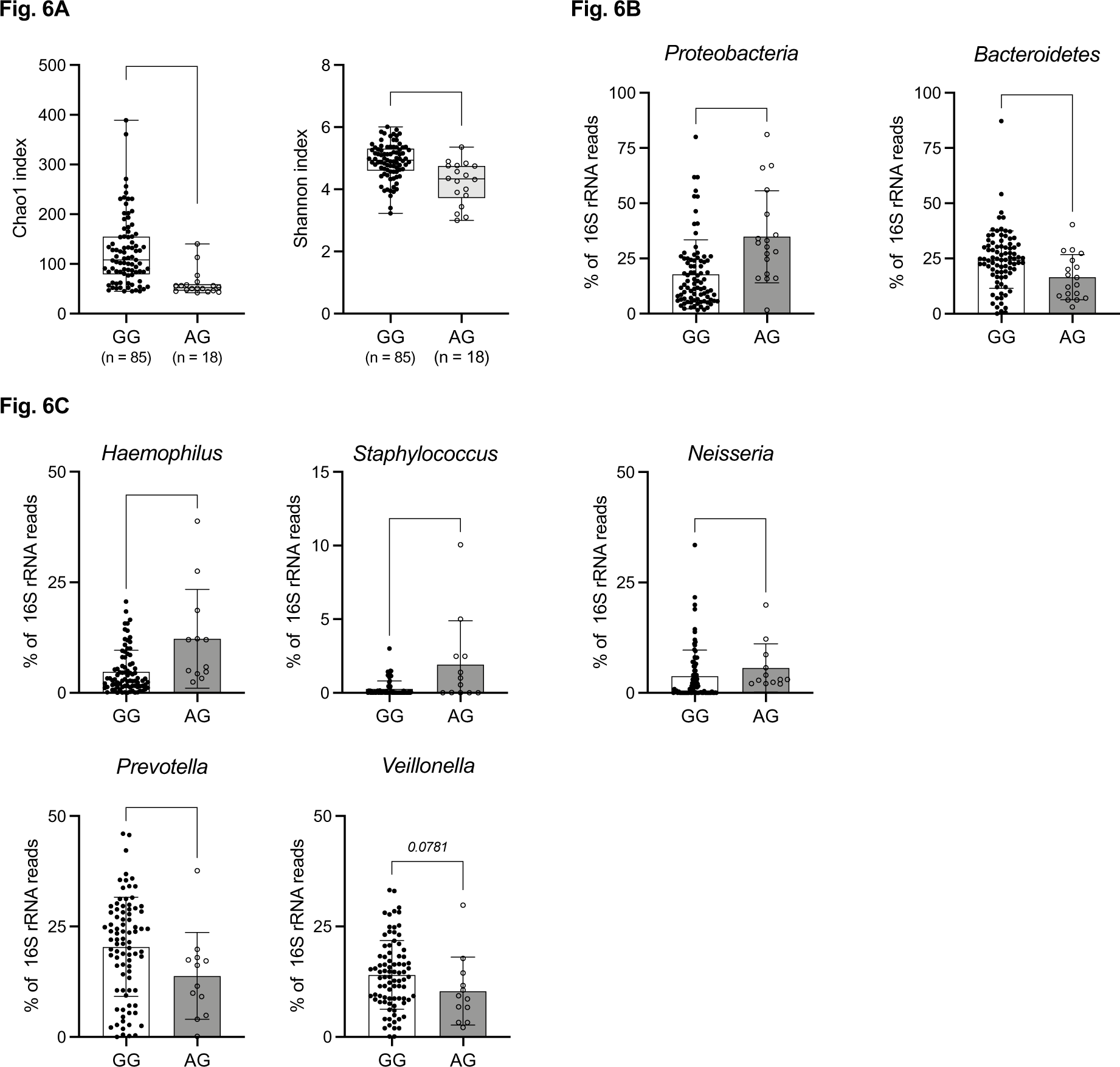
*TLR5* rs5744168 modifies lung microbiome and TLR5 effect on fibrosis depends on the presence of microbiome. **(A)** left: Chao1 index and right: Shannon Diversity Index of BALF microbiota of rs5744168 minor allele non-carrier (N = 85) and carrier (N = 18) IPF patients. **(B)** Phylum composition of Proteobacteria, Bacteroides, Firmicutes, and Actinobacteria of BALF microbiota of rs5744168 minor allele non-carrier and carrier IPF patients. **(C)** Genus composition of Haemophilus, Staphylococcus, Neisseria, Prevotella, and Veillonella of BALF microbiota of rs5744168 minor allele non-carrier and carrier IPF patients. P-values from ANOVA with Bonferroni post-hoc correction.

## DISCUSSION

Recent research has shed considerable insight into the pathogenesis of IPF, highlighting innate host defense among the emerging disease pathways. The airway mucin gene *MUC5B* is the most significant IPF genetic risk factor (*19*), while genes in the toll-like receptor pathway may influence either susceptibility or disease progression (*3, 20–22*). There is also accumulating evidence that the microbiome mediates fibrosis development ^5, 6, 21,25, 27^. In mice, lung dysbiosis precedes lung fibrosis, while microbiome eradication ameliorates fibrosis (*5, 7*). In IPF patients, lung dysbiosis is associated with IPF progression and decreased survival (*4–6*). The overriding principle connecting genetic and microbiome observations may be that dysfunctional host defense promotes dysbiosis, which perpetuates a chronic inflammatory status that perturbs lung homeostasis (*23*). Here, we report a novel genetic risk factor for IPF, *TLR5* SNP rs5744168, and demonstrate that TLR5 dysfunction promotes fibrosis because it confers failure to control lung dysbiosis. Microbial genera, associated with disease progression, are also associated with activation of immune pathways in IPF patients (*24*), supporting the experimental finding that dysbiosis may promote fibrosis through immune activation (*7*). Our findings suggest that the immunity-microbiome interaction in fibrosis is bidirectional: while dysbiosis promotes *aberrant* immune activation, *Tlr5* activation ameliorates dysbiosis and fibrosis through a *homeostatic* immune activation.

Tlr5 activation has antibiotic properties in various lung infection models, including flagellated, non-flagellated, gram-positive and -negative bacteria (*25–27*). We now show that TLR5 activation modulates the microbiome in non-infectious lung injury and that TLR5 protects against fibrosis by reducing dysbiosis in the injured lung. It is notable that flagellin-induced activation pathways (*e.g*. NLRC4, CAMP) are positively associated with lung dysbiosis and blood neutrophilia in IPF patients (*24*). This may suggest that flagellated bacteria play a role in IPF pathogenesis, perhaps by inducing a feedback loop which modulates lung microbiota. We found that, in IPF patients, TLR5 deficiency is associated with an increase in Proteobacteria, which include several flagellated genera (*e.g. Pseudomonas, Legionella*); these may activate lung TLR5 and induce a feedback loop suppressing them. In the mouse gut, *Tlr5* deficiency promotes an unstable microbiome, also with preponderance of Proteobacteria, which promotes chronic inflammation (*28*). We postulate that similar effects of TLR5 deficiency may prevail in the lung and induce chronic inflammation conducive to fibrosis development after injury.

In our study, epithelial TLR5 activation mediates the protective effect of the TLR5 agonist. This mirrors findings in the mouse gut, where epithelial Tlr5 activation shapes microbial composition (*9*). Our findings support the central role of epithelia in lung homeostasis and suggest that epithelia help regulate lung microbiome. Notably, smoking suppresses airway epithelial *TLR5* expression and responsiveness to flagellin in humans (*29*). Such acquired TLR5 deficiency may exemplify how environmental exposures precipitate lung injury.

In our analysis, the rs5744168 minor allele had a clear association with IPF in homozygotes of the *MUC5B* major allele but not in *MUC5B* minor allele carriers. This may simply reflect the dominant role of MUC5B on IPF susceptibility. However, it may also suggest intersecting pathological mechanisms. In mice, muc5b is required for lung microbial clearance (*30*), while in IPF patients *MUC5B* polymorphism is associated with microbial burden (*6*), supporting the host immunity-microbiome relationship that we also demonstrate. It is conceivable that *MUC5B* overexpression, induced by the rs35705950 minor allele, alters the microbiome in ways that cannot be impacted by TLR5.

Our results support a role for the lung microbiome in IPF pathogenesis. Eradication of lung microbiome ameliorates lung fibrosis while eradication of gut microbiome does not (*7*). We demonstrate that airway epithelial Tlr5 mediates the TLR5-agonist effect on lung fibrosis and show that lung *Tlr5* expression is altered in the presence of microbiome. In aggregate, our data support that airway epithelial TLR5 modulation of the lung microbiome ameliorates lung fibrosis. TLR5 deficiency did not impact microbial burden in IPF patients, however it did alter lung microbial composition. While microbial burden is associated with IPF progression (*5, 6, 31*), it may simply be a sensitive indicator of dysbiosis. Bacterial burden did not show an association with BALF cytokines in IPF patients, whereas dysbiosis did; furthermore, antibiotics ameliorate bleomycin-induced fibrosis by reducing dysbiosis without changing total microbial burden (*5*). This supports that dysbiosis, more than microbial abundance, may activate lung immunity and disrupt epithelial homeostasis. By the same token, immune pathways that prevent dysbiosis, like TLR5 activation, may ameliorate fibrosis without an obvious effect on total microbial burden.

In conclusion, our results demonstrate that TLR5 deficiency is a risk factor for IPF, and that TLR5 activation ameliorates fibrosis by reducing dysbiosis. TLR5 may thus be a potential IPF therapeutic or prophylactic target. More generally, future preventative measures against IPF may involve modulation of the lung microbiome in select patients, perhaps those with evidence for dysbiosis and aberrant immune activation. Conversely, the homeostatic activation of the immune system by microbial derivatives may be a novel modality to impact development or exacerbation of lung disease without need for chronic antibiotic treatment and its associated side effects.

## MATERIALS AND METHODS

### Human subject approval

All human studies were approved by the respective Institutional Review Boards. (University of Pittsburgh, IRB# STUDY19040326 and STUDY20030223, University of Colorado COMIRB 15-1147). Written informed consent was obtained for all subjects before enrollment in the research study. For the discovery cohort, IPF subjects were recruited from the Simmons Center for Interstitial Lung Diseases at the University of Pittsburgh Medical Center. Diagnosis of IPF was supported by clinical, physiologic, and high-resolution computed tomography studies (*32*). Patients fulfilled the criteria of the American Thoracic Society and European Respiratory Society for the diagnosis of IPF (*33, 34*). Patients with known causes of interstitial lung disease were excluded. Unrelated healthy subjects with no self-reported advanced lung diseases were recruited from the University of Pittsburgh Medical Center with written informed consents (STUDY19070274) and donors with discarded donated blood recruited from the Great Pittsburgh Blood Bank under an exempt human subject study (PRO11080146) were used as controls. University of Colorado replication cohort consisted of subjects consented for genetic studies of IPF approved by the University of Colorado Combined Institutional Review Board (COMIRB 15-1147). IPF diagnosis was established using the same ATS/ERS definitions of IPF as the discovery cohort. We excluded cases with known causes for interstitial lung disease. Control subjects were consented through COPDGene and approval for genotyping for this study was obtained through COMIRB 15-1147. Approval for the study at Imperial College was obtained from the local research ethics committee (15/SC/0101 and 15-LO-1399).

### Genotyping and statistical analysis of genetic association

Genotyping of *TLR5* SNP rs5744168 and *MUC5B* SNP rs35705950 were performed on DNA extracted from whole blood (PAXgene Blood DNA Kit, Qiagen, Germantown, MD) using Taqman SNP genotyping assay C 25608804_10 and C 1582254_20, respectively (Thermo Fisher, Waltham, MA). 20 ng of DNA were included in each reaction which were run on a ViiA7 Real-Time PCR machine per the SNP genotyping assay instructions. We examined the target TLR5 SNP rs5744168, as well as the established IPF risk factor MUC5B SNP rs35705950 as a positive control for observed IPF associations, in both the discovery dataset (Pittsburgh) as well as in the validation dataset (Denver). We then conducted a meta-analysis combining results from the two datasets. SNPs were tested for satisfaction of the Hardy-Weinberg Equilibrium (HWE) assumption. Based on previous meta-analytic evidence, we employed additive modeling for rs35705950; we tested both additive and dominant models for rs5744168 as suggested by Hawn et al. (*12, 14, 54*). Alternative analyses tested the models for each SNP, combining the case data from Pittsburgh and Denver, and using unscreened controls (non-Finnish European) from the GNOMAD database (https://gnomad.broadinstitute.org/). PLINK version 1.07 and SAS version 9.4 were used for analyses, and METAL was used for meta-analysis. To test for the interaction of the two SNPs, we combined the discovery and validation datasets to maximize power and tested for interaction of the SNPs using logistic regression (additive rs35705950 interacting with additive rs5744168, and additive rs35705950 interacting with dominant rs5744168).

### Analysis of human scRNA-Seq data

Publicly available single cell RNA-Seq (scRNA-Seq) data from Reyfman *et al*. (*35*) (GSE122960) and Adams *et al*. (*36*) (GSE136831) (also available through www.IPFCellAtlas) were analyzed for *TLR5* gene expression. First, count matrices were initialized using the R (R4.0.2) package “Seurat” (*37*). Count matrices were then filtered to remove cells containing low (<200) or high (>5,500) gene content, low (<250) or high counts (>25,000), low (0.1%) or high (>15%) mitochondrial content, and low (<0.1%) or high (>20%) cycling cells. The combined count matrices containing only high-quality cells were then used for further analyses. Filtered count matrices from both sources (Reyfman *et al.* & Adams *et al*.) were integrated using the “Harmony” (*38*) package. Using this algorithm, we were able to simultaneously adjust any experimental and biological factors by projecting cells into a shared principal component space for embedding, allowing for combined analyses of these two large scRNA-Seq datasets. In sum, known co-variates (e.g. sex, age, disease status) and unknown co-variates were corrected for, allowing analysis of same cell types across any co-variates. Clustering of scRNA-Seq data was performed using the Leiden community detection algorithm (*39*) from the “monocle3” (*40*) R package using R3.6.1. Using resolution of 0.00025, we were able to distinguish different cell types as annotated by Adams *et al*. (*36*) using Uniform Manifold Approximation and Projection (UMAP) dimensions as input. Cell types constituting clusters were identified using previously annotated metadata from Adams *et al*. (*36*). In addition, several key markers were used to validate the previously annotated cell types. In some instances (*e.g.* alveolar macrophages) clusters were merged according to cell type markers and previous annotations. These clusters were projected onto UMAP space for visualization. TLR gene expression was summarized within each major annotated cell type and visualized using dot plots and stratified by disease status.

### Experimental models of lung fibrosis

The following strains were purchased from Jackson Laboratory (Bar Harbor, ME): *Tlr5^Fx/Fx^* (B6(Cg)-*Tlr5^tm1.1Gewr^*/J), *LysM-Cre* (B6.129P2-*Lyz2^tm1(cre)Ifo^*/J), *Trif^-/-^*(B6.129P2-Ticam<TM1AKI>), *Myd88^-/-^* (B6.129P2(SJL)-Myd88<tm1.1Defr>/J), and *Tlr5^-/-^* (B6.129S1-Tlr5<TM1FLV>/J). *SPC-Cre* (B6;D2-Tg(Sftpc-cre)1Blh) (*41*) and *Rosa-Tomato Spc-CreER* (B6;Cg-Gt(ROSA)26Sor<tm9(CAG-tdTomato)Hze> Sftpc<tm1(cre/ERT2)Blh>) (*42*) mice were graciously provided by Brigid Hogan (Duke University) and Donald N. Cook (NIEHS/NIH). *Tlr5* lung epithelium (*Tlr5^Fx/Fx^ Spc-Cre*) and myeloid (*Tlr5^Fx/Fx^ LysM-Cre*) specific knockouts were generated by crossing *Tlr5^Fx/Fx^* mice to *Spc-Cre* and *LysM-Cre* mice, respectively. All mice were bred and maintained at the NIEHS/NIH animal breeding facility under specific pathogen-free conditions. All animals were single-housed in static cages after weaning (to avoid cage effects on microbiome analysis), were kept in a 12-hour dark-light cycle, and were provided sterilized chow and water, *ad libitum*. All animal studies were approved and conducted in accordance with the NIEHS/NIH guidelines for the humane treatment of animals. All studies were approved by the NIEHS Animal Care and Use Committee (ACUC).

#### Bleomycin model

Mice were anesthetized using isoflurane or ketamine (100 mg/kg)/xylazine (7 mg/kg) prior to bleomycin sulfate (Sigma, B8416) administration by oropharyngeal aspiration or by intratracheal instillation, respectively, at the indicated concentrations in 50 μL normal saline. Animals that received ketamine/xylazine were administered atipamezole (0.5 mg/mL x 0.1 mL/mouse) for the reversal of the sedative and analgesic effects. Animals were euthanized on days 5-7 for evaluation of epithelial injury and microbiome analysis, and on days 14-21 for evaluation of fibrosis.

#### TLR5-agonism in bleomycin model

TLR5-agonists, derived from Salmonella flagellin (*16, 43*), were provided by Genome Protection, Inc. Mice were dosed with 1.0-1.5 μg of TLR5-agonist subcutaneously (200 μL volume in normal saline with 0.01% Tween-80), 30-minutes prior to bleomycin administration or at indicated timepoints.

#### Antibiotics in bleomycin model

Animals were given a mixture of Ampicillin (1 g/L, Sigma, A1593-25G), Vancomycin (500 mg/L, Sigma, V2002-5G), Neomycin Sulfate (1 g/L, Sigma, N5286-25G), and Metronidazole (1 g/L, M1547-25G) in drinking water and in mash for 4-weeks prior to experimental use. Sucralose (1/50 dilution) was added for palatability.

### Histology and immunohistochemistry

Lungs were inflated with 4% paraformaldehyde, sectioned at 5 μm and stained with hematoxylin and eosin or Masson’s trichrome.

#### Immunofluorescence staining of human IPF lung

Formalin-fixed, paraffin embedded sections of human IPF lung were purchased from the Duke Biospecimen Repository & Processing Core. Sections were deparaffinized in xylene and rehydrated through graded ethanol to Tris Buffered Saline with 1% Tween 20 (1XTBST). Following 10 minutes in 3% aqueous hydrogen peroxide, samples were subjected to antigen retrieval in 1X EDTA antigen retrieval solution (Fisher Scientific) in a pressure cooker (Biocare Medical, Pacheco, CA) for 5 minutes. Tissues were blocked in 10% Normal Donkey Serum (Jackson ImmunoResearch, West Grove, PA), 1% BSA, in 1XTBST for one hour at RT, then incubated with primary antibodies overnight at 4°C. Antibodies were rabbit anti-Tlr5 (Origene, Rockville, MD; 1:200), and mouse anti-SP-C (Santa Cruz, Dallas, TX; 1:100). Secondary antibodies (Donkey anti-mouse Alexa Fluor 594, 1: 200, and Donkey anti-rabbit Alexa Fluor 488; 1:100; both from Invitrogen) were applied for 1 hour at room temperature. After washing in 1XPBS, samples were incubated with DAPI solution (ThermoFisher Scientific) for 15 minutes, washed with 1XPBS and cover-slipped with Prolong Diamond Antifade (ThermoFisher). Images were acquired using a Zeiss LSM 780 inverted confocal microscope.

#### Immunofluorescent staining of murine lung

Lung samples were fixed in 4%formaldehyde in PBS overnight at +4oC, washed with PBS, cryopreserved in 15%-30% sucrose/PBS and embedded in Neg-50 frozen section medium (Fischer Scientific). 12-thick cryosections were stained in cocktail of rabbit monoclonal antibody against NFκB p65 (Cell Signaling, #8242; dilution 1:200) and rat monoclonal anti-cytokeratin 8 Troma-I antibody (DSHB; dilution 1:200) for 1 hour at room temperature. Secondary antibodies were donkey anti-rabbit IgG AlexaFluor488-conjugated and donkey anti-rat IgG AlexaFluor546-conjugated (2μg/ml each, both from Jackson ImmunoResearch). Sections were mounted with ProLong Gold or anti-fade reagent with DAPI (Invitrogen). Images were captured using Zeiss Axio Imager Z1 microscope equipped with epifluorescent light source and AxioCam MR3 camera and processed with AxioVision 4.6 software.

### Hydroxyproline assay

Whole lungs were processed for this assay. Lung hydroxyproline contents were analyzed following the Hydroxyproline Assay Kit (Catalog #: QZBTOTCOL5) from QuickZyme Biosciences. Data are expressed as micrograms of hydroxyproline per lung.

### Preparation and flow sorting of epithelial cells

*Rosa-Tomato Spc-CreER* (B6;Cg-Gt(ROSA)26Sor<tm9(CAG-tdTomato)Hze> Sftpc<tm1(cre/ERT2)Blh>) and *Tlr5^Fx/Fx^ SPC-Cre* (*B6.Cg-Tlr5<tm1.1Gewr> Tg(Sftpc-cre)1Blh Tg(Vil1-cre)1000Gum*) mice were utilized for this experiment. Rosa-Tomato SPC-CreER mice were injected with Tamoxifen (0.25 mg/10 g to 2.5 mg/10 g body weight) as previously described (*44*). After euthanasia with FatalPlus, lungs were perfused and inflated with digestion media containing Elastase (4.5 U/mL, Worthington, Catalog #: LS002280) and DNase (20 mg/mL, Sigma, Catalog #: DN25-1G) for a total of 1 hour. Lungs were then minced, and single cells were isolated using a 70 μm filter. Type 2 alveolar epithelial cells were sorted for EpCAM^+^ TOMATO^+^ by flow cytometry. We also analyzed EpCAM^+^ TOMATO^-^ cells, which include bronchial epithelial cells and some Type 1 alveolar epithelial cells. *Tlr5^Fx/Fx^ SPC-Cre* mice were euthanized with FatalPlus and lungs were collected, perfused, and digested as described above. Type 2 alveolar epithelial cells were sorted as described by Nakano *et al*. (53).

### Analysis of murine RNA-Seq data

#### RNA-seq data analysis

Flow-sorted cells were subjected to RNAseq analysis. Raw reads were filtered by a custom perl script to remove poor quality reads (average quality score < 20), and then the cutadapt program (*45*) (v2.1) was applied to remove adaptor sequences. The preprocessed reads were aligned to the mouse mm10 genome using the Spliced Transcripts Alignment to a Reference (STAR) aligner (*46*) (version 2.7.0f). Subread featureCounts (version 1.4.6) was applied for gene quantification based on University of California Santa Cruz (UCSC) mm10 RefSeq gene annotation. The raw count matrix was used to detect the differential gene expression between treatment and the control samples by DESeq2 package (version 1.22.2). Differentially expressed genes were determined using the following cutoffs: (1) mean FPKM ≥1 in at least one of the conditions; (2) fold change over 1.5; and (3) BH-adjusted p-value < 0.05. The RNAseq data was deposited to NCBI Gene Expression Omnibus repository (GEO accession number GSE166852). *Gene Set Enrichment Analysis*. Gene Set Enrichment Analysis (GSEA) was performed using the GSEA software (version 4.0.0) with the Molecular Signatures Database (*47*) (MSigDB, version 7.0). The preranked gene list for each comparison was sorted based on the score generated from log2FoldChange multiplied by negative log10 p-value. The “GSEAPreranked” tool was performed with the classic scoring scheme and 1000 times of permutation.

### Murine Lung Microbiome Analysis

Animals were euthanized using FatalPlus by intraperitoneal administration. Lungs were excised from animals under sterile conditions and flash frozen in liquid nitrogen for storage (-80 °C) until sample processing.

#### 16S Ribosomal RNA gene sequencing

DNA was isolated, the V4 region of the bacterial rRNA gene amplified and sequenced using previously published protocols that employ multiple DNA isolation, procedural and sequencing controls to control for the effects of contamination in low biomass specimens (*5, 48*). The MiSeq platform was used for 16S community sequencing (Illumina).

#### Bacterial DNA measurement

Bacterial DNA was quantified in experimental specimens and isolation and procedural controls using a previously published protocol (*48*) with a QX200 droplet digital PCR (ddPCR) system (Bio-Rad).

#### Statistical analysis

16S sequence data was processed and analyzed using mothur version 1.44.3 software (*49*). The vegan package was used to further analyze community sequencing data. All analysis was carried out in R version 4.0.0. (R Core Team (2020). R: A language and environment for statistical computing. R Foundation for Statistical Computing, Vienna, Austria (https://www.R-project.org/).

### IPF patient bacterial DNA isolation and quantification

Genomic DNA was extracted from bronchoalveolar lavage fluid (BALF) cell pellets, as previously described (*31*). Aliquots of 2 ml of the BAL samples were centrifuged (21,000 x *g*; 30 minutes) to pellet cell debris and bacteria. Pellets were transferred to lysing matrix E (LME) tubes (MP Biomedicals, USA) and resuspended in 100 μl of supernatant, to which 500 μl acetyl trimethylammonium bromide (CTAB) buffer (10% w/v CTAB in 0.5 M phosphate buffered NaCl) and 500 μl phenol-chloroform-isoamyl alcohol (25:24:1) were added, and shaken (5.5 m s^-1^; 60 seconds) in a FastPrep Instrument (MP Biomedicals, USA). Samples were then extracted with an equal volume of chloroform-isoamyl alcohol (24:1), and DNA precipitated with 2 volumes of precipitation solution (30% w/v PEG6000 in NaCl). Pellets were resuspended in 100 μl Tris-EDTA following ethanol washing. The quality and quantity of the extracted DNA was measured using a NanoDrop™ 1000 Spectrophotometer (Thermo Fisher Scientific, UK) and the DNA was stored at -80 °C until further analysis.

#### 16S rRNA sequencing and bioinformatics analysis

Paired-end sequencing of the hypervariable V4 region of the 16S rRNA gene was performed with the 515 forward and 806 reverse primers using an Illumina MiSeq instrument (Illumina, San Diego, CA) to produce 150 base-paired reads. Bioinformatics analysis of bacterial 16S rRNA amplicon data was conducted using the Quantitative Insights Into Microbial Ecology (QIIME2, version 2019.10) pipeline (*50*) as previously described (*51*). Downstream pre-processing was performed in R (version 3.4.3) to obtain the relative abundances of the microbiota in each sample and to visualize the Shannon and Chao1 α-diversity indexes.

#### 16S rRNA gene quantitative polymerase chain reaction

Triplicate 10 μl quantitative polymerase chain reactions (qPCR) were set up containing 1 μl of bacterial DNA and 9 μl of Femto™ bacterial qPCR premix (Cambridge bioscience, UK) as previously described (*31*).

## Supplementary Materials

### Supplementary Figures

Fig. S1. UMAP of human lung cells.

Fig. S2. Immunohistochemical staining of human lung sections with TLR5 blocking peptide.

Fig. S3. Immunohistochemical staining of human IPF lung sections with TLR5 blocking peptide.

Fig. S4. Tlr5 protects against bleomycin-induced lung injury and fibrosis.

Fig. S5. Lung *Tlr5* expression after intranasal flagellin treatment and in germ-free mice.

Fig. S6. Experimental setup for microbiome reconstitution.

Fig. S7. TLR5 polymorphism is associated with dysbiosis in IPF patients.

### Supplementary Tables

Table S1. Characteristics of cases and controls.

Table S2. rs5744168 and rs35705950 genotypes by study source and case/control status.

Table S3. Combined Discovery and Validation Genotypes.

Table S4. Associations with IPF, for rs5744168 (additive and dominant) and rs35705950 (additive) using combined cases from discovery and validation samples, and unscreened non-Finnish European controls from gnomAD.

Table S5. Combined discovery and validation data.

Table S6. Logistic regression with interaction of additive rs5744168 and additive rs35705950.

Table S7. Logistic regression with interaction of dominant rs5744168 and additive rs35705950.

Table S8. Top 100 enriched gene sets in AEC2s in response to TLR5-agonist.

Table S9. Top 100 enriched gene sets in bronchial epithelial cells in response to TLR5-agonist.

Table S10. *Tlr5*-deficient lung epithelia fail to upregulate antimicrobial factors in response to TLR5-agonism.

## Author contributions

Conceptualization: SG, AVG

Investigation: YS, EW, AS, CGJ, XM, SH, SM, CST, SYT

Statistical analysis of the genetic associations: JMcG

Human genotyping and gene expression data generation and analysis: PK, KG, YZ, JHL, KG, AW, IY, NK, DAS

Mouse microbiome analysis: JHL, DNo’D

Mouse RNA seq analysis: LZ and JLL

Human microbiome data analysis: PLM and RI

Writing – original draft: YS, SG

Writing – review and editing: All authors.

### Data and materials availability

All data are available in the main text or the supplementary materials. Gene expression analysis and microbiome analysis data are deposited in public databases and the respective accession numbers are entered in the Materials and Methods section.

