## supplemental data for "Toll-Like-Receptor 5 protects against pulmonary fibrosis by reducing lung dysbiosis"

#### Supplemental Methods

Case-control genotypes in the cohorts

In the validation dataset both rs5744168 and rs35705950 met HWE ($p = 0.8523$ and 0.003 respectively). In the combined dataset both SNPs met HWE. Comparing cohorts, the METAL heterogeneity test p-value for rs35705950 was significant ($p = 0.008$) but was non-significant for rs5744168 ($p = 0.742$). The heterogeneity and discovery cohort HWE results probably reflect the relatively small numbers of the discovery cohort. Similar results were obtained using the PLINK meta-analysis module. **Supplementary Table 2** provides the genotype breakdown by sample source and SNP, and **Supplementary Table 3** provides the genotype breakdown after combining samples, for subjects with complete genotypes for rs5744168 and rs35705950 (N = 3867).

Alternative analysis with unscreened controls

We performed an alternative analysis using an additional set of control subjects, to further confirm that our results are generalizable. The gnomAD (genome aggregation database) provides variant frequencies based on aggregation of exomic and genomic data from unrelated individuals screened in disease-specific and population studies [1]. We obtained the non-Finnish European data for rs5744168 and rs35705950 (accessed March 28, 2019) and calculated “control” allele frequencies for comparison with our combined case data (**Supplementary Table 4**). The additive model ORs [95% CI] for IPF were 1.27 [1.09 – 1.49, $p = 0.0028$] and 4.96 [4.41 – 5.59, $p < 0.0001$] for rs5744168 and rs35705950, respectively, consistent with the results we obtained in our discovery and validation cohorts.

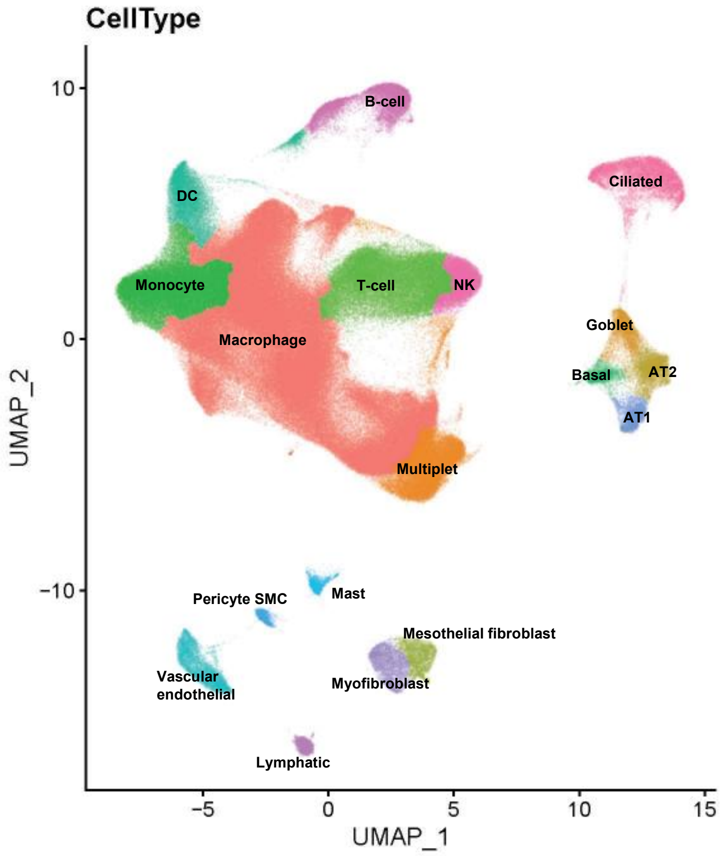

**Supplementary Figure S1. UMAP of human lung cells** from 36 control, 18 COPD, 1 HP, 36 IPF, and 2 SScILD lungs, labeled by cell type, from [2] and [3].

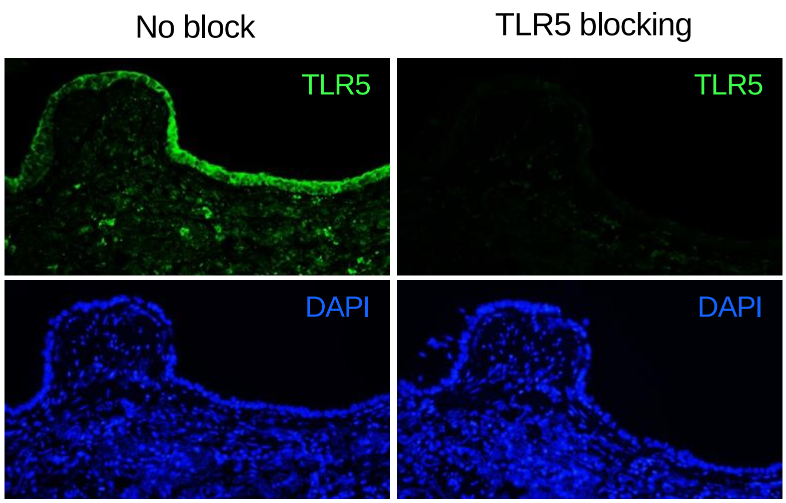

**Supplementary Figure S2. Immunohistochemical staining of human lung sections with TLR5 blocking peptide** (right) and without TLR5 blocking peptide (left); green: TLR5, blue: DAPI. Representative images shown.

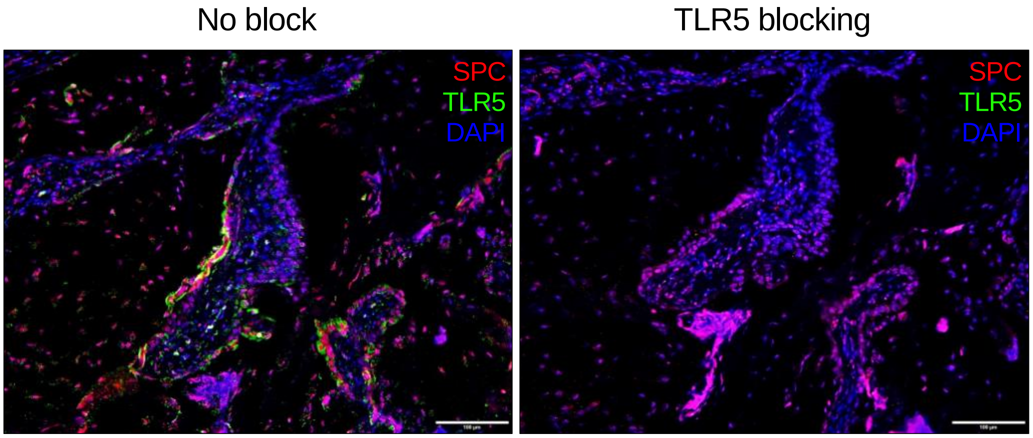

**Supplementary Figure S3. Immunohistochemical staining of human IPF lung sections with TLR5 blocking peptide** (right) and without TLR5 blocking peptide (left); red: SPC, green: TLR5, blue: DAPI. Representative images shown.

**Supplementary Fig. 4A**

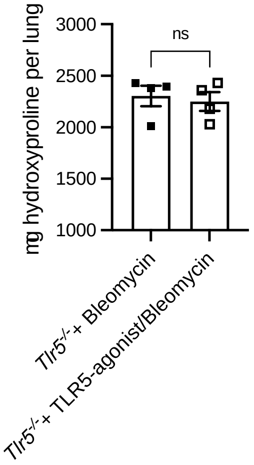

**Supplementary Fig. 4B**

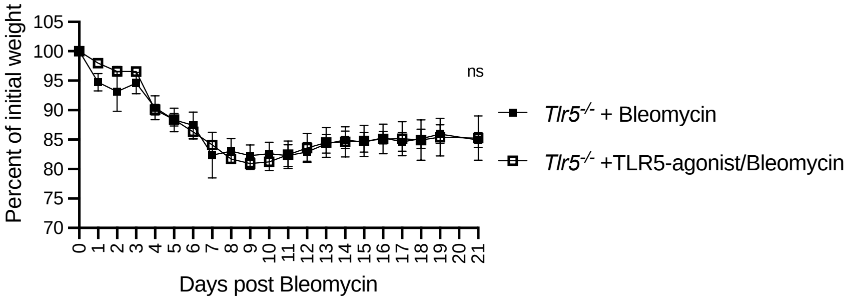

**Supplementary Fig. 4C**

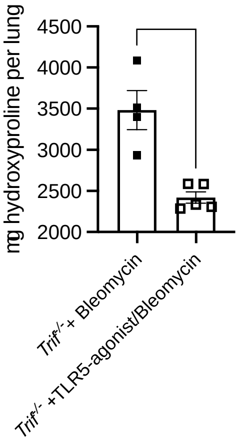

**Supplementary Fig. 4D**

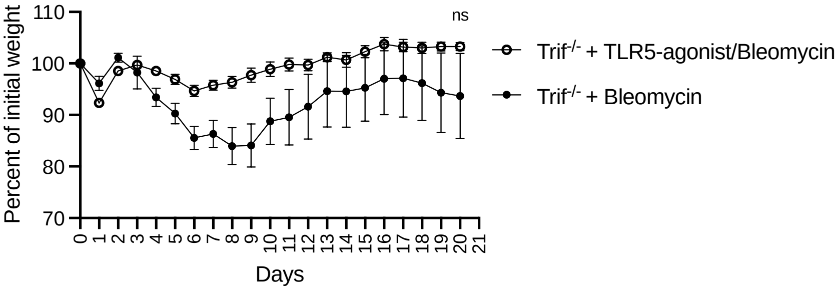

**Supplementary Fig. 4E**

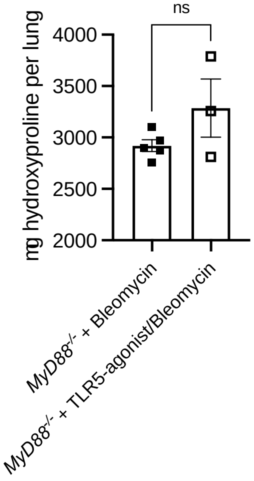

**Supplementary Fig. 4F**

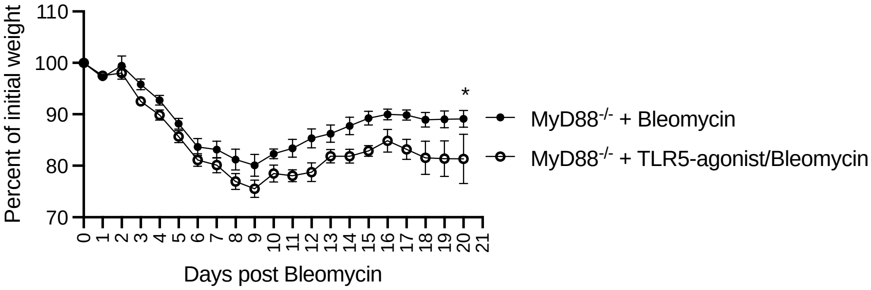

**Supplementary Fig. 4G**

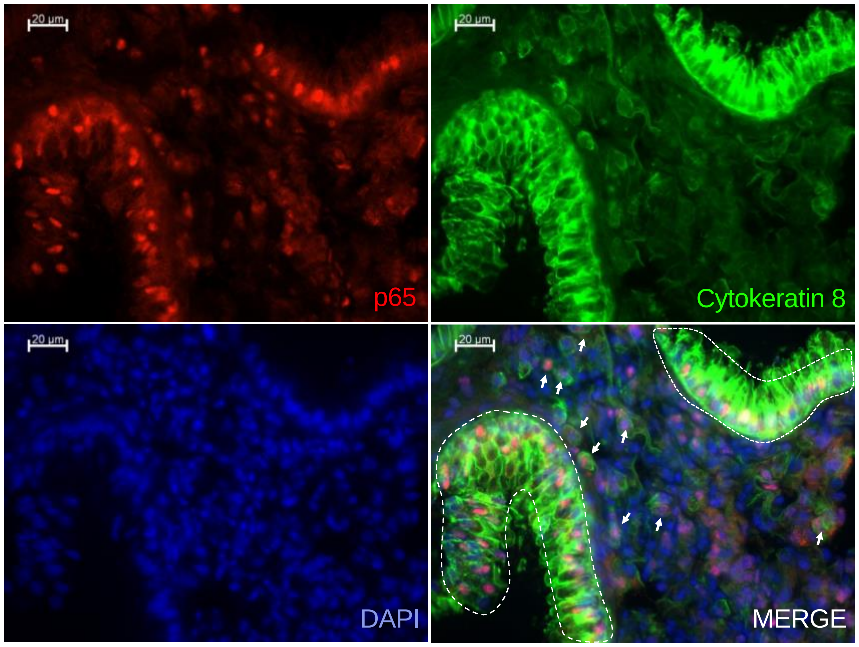

**Supplementary Fig. 4H**

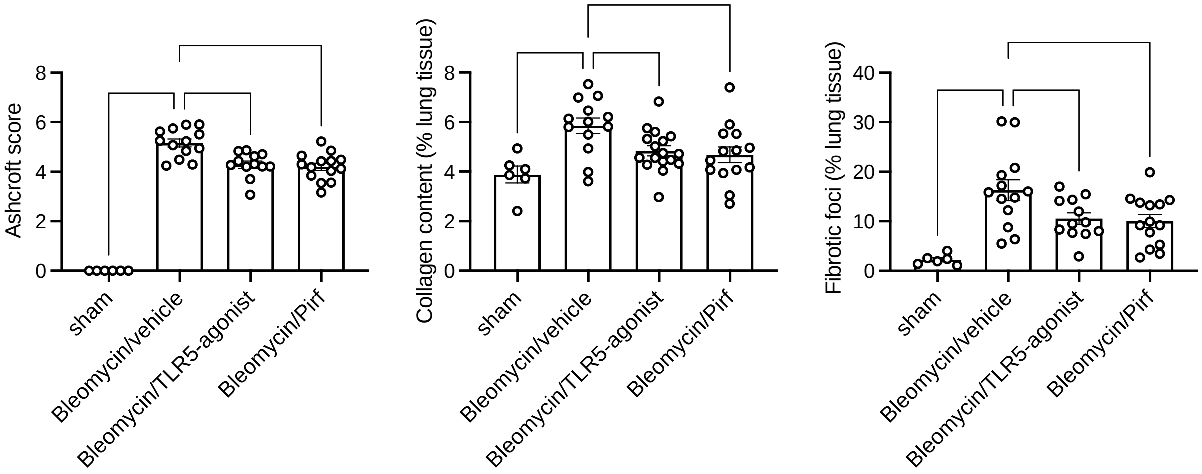

**Supplementary Figure S4. TLR5 protects against bleomycin-induced lung injury and fibrosis.**

**(A)** Quantification of total collagen per lung determined by hydroxyproline assay, 21 days post bleomycin administration in vehicle and TLR5-agonist treated *Tlr5*$^{-/-}$ mice. Experiment repeated over 5 times. **(B)** Percent of initial weight of vehicle or TLR5-agonist treated *Tlr5*$^{-/-}$ animals over

21 days post bleomycin administration. (**C)** Quantification of total collagen per lung determined by hydroxyproline assay, 21 days post bleomycin administration in vehicle and TLR5-agonist treated *Trif*$^{-/-}$ mice. (**D)** Percent of initial weight of vehicle or TLR5-agonist treated *Trif*$^{-/-}$ animals over 21 days post bleomycin administration. (**E)** Quantification of total collagen per lung determined by hydroxyproline assay, 21 days post bleomycin administration in vehicle and TLR5-agonist treated *MyD88*$^{-/-}$ mice. (**F)** Percent of initial weight of vehicle or TLR5-agonist treated *MyD88*$^{-/-}$ animals over 21 days post bleomycin administration. **(G)** Immunohistochemical staining of lung sections TLR5-agonist treated mice, 2 hours post treatment; red: p65, green: cytokeratin 8, blue: DAPI, white: merge. White arrows and discontinuous line surrounding airways indicate Cytokeratin 8 and p65 double positive cells. Representative images shown. **(H)** Vehicle, TLR5-agonist (0.73μg/mouse), or Pirfenidone (Pirf, 100mg/kg) were given at various time points after bleomycin (2mg/kg) administration (Day 0). Vehicle and TLR5-agonist were given at Days 7, 11, 14, and 18 post bleomycin administration. Pirfenidone was given daily starting Day 7 post bleomycin administration. Left: Ashcroft score; center: Collagen Content (% lung tissue); and right: Pulmonary Foci (%). Analysis was performed with ANOVA with Holm-Sidak correction *p<0.05, ** p<0.01, *** p<0.001, **** p<0.0001.

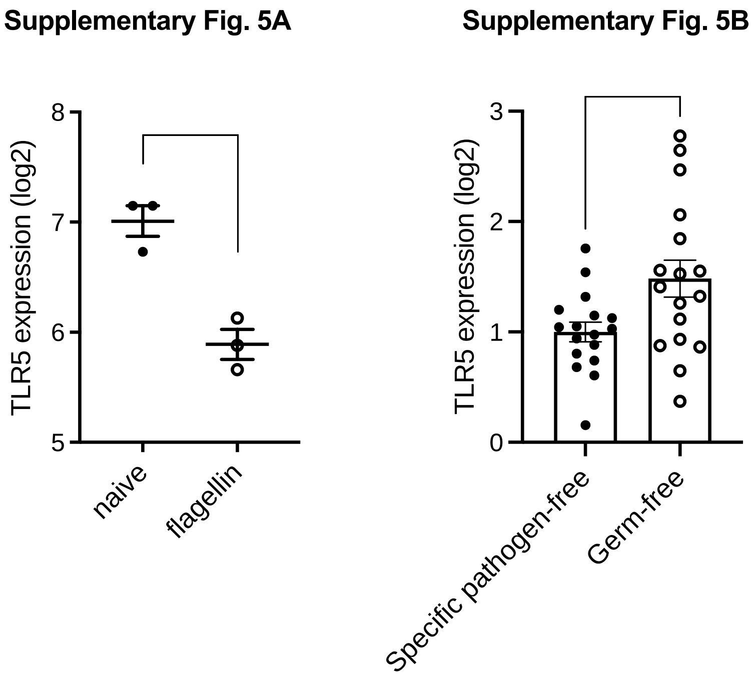

**Supplementary Figure S5. Lung *Tlr5* expression after intranasal flagellin treatment and in germ-free mice.**

**(A)** Mice were exposed by oropharyngeal aspiration to 50 µL PBS containing 100 µg recombinant flagellin from S. typhimurium (Cat# tlrl-flic-50, InvivoGen) daily for 3 days, and lungs were collected on day 4 for *Tlr5* gene expression, determined by quantitative PCR. **(B)** *Tlr5* gene expression of lungs from specific pathogen (SPF) or germ-free (GF) conditions, determined by quantitative PCR.

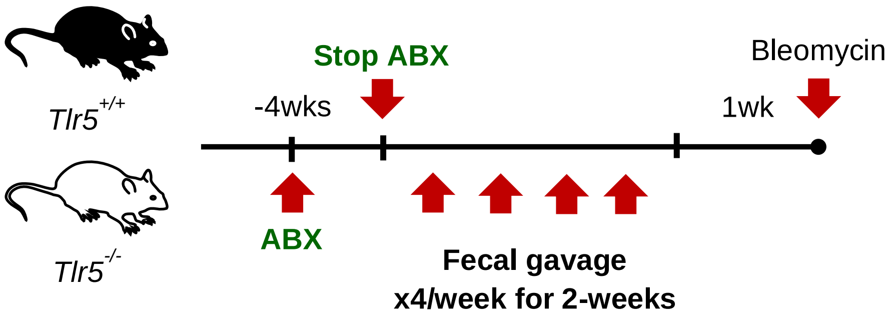

**Supplementary Figure S6. Experimental setup for microbiome reconstitution.**

**(A)** Animals were treated with an antibiotic cocktail for 4-weeks prior to microbiome reconstitution. Fecal gavages were given x4/week for 2-weeks, followed by a 1-week rest period prior to bleomycin-administration.

Supplementary Fig. 7

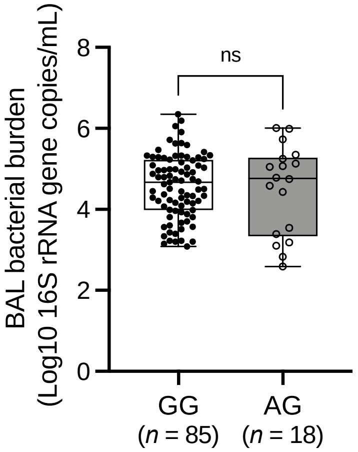

**Supplementary Figure S7. TLR5 polymorphism is associated with dysbiosis in IPF patients.** BALF bacterial burden, determined by 16s quantification, of rs5744168 minor allele non-carrier and carrier IPF patients.

**Supplementary Table S1. Characteristics of cases and controls.**

| | | Discovery Sample | | Validation Sample | |
|---|---|---|---|---|---|
| | | Pittsburgh cases | Pittsburgh controls | Denver cases | COPD controls |
| N | rs5744168 | 277 | 397 | 833 | 2,505 |
| | rs35705950 | 275 | 386 | 697 | 2,500 |
| Female (%) | rs5744168 | 32.1 | 55.4 | 30.0 | 50.6 |
| | rs35705950 | 31.6 | 55.4 | 24.0 | 50.6 |
| Age (SD) | | 68.1 (8.9)[a] | 48.1 (15.2)[b] | 65.4 (11.0)[c] | 59.5 (8.7)[d] |
| FVC % predicted | | 64.0 (18.9)[e] | - | 69.0 (19.1)[f] | - |
| DLCO % predicted | | 43.0 (15.9)[g] | - | 52.7 (21.5)[h] | - |
| Ever smoked (%) | | 70.4 [i] | 35.7 [j] | 72.7 [k] | 100.0 [l] |
| [a] N=277; [b] N=397; [c] N=769; [d] N=2500; [e] N=246; [f] N=415; [g] N=191; [h] N=533; [i] N=277; [j] N=210; [k] N=686; [l] N=2505 | | | | | |

**Supplementary Table S2. rs5744168 and rs35705950 genotypes** by study source and case/control status.

| Source | Status | rs5744168 | N | rs35705950 | N |
|---|---|---|---|---|---|
| Discovery | control | AA | 10 | TT | 4 |
| | | AG | 32 | GT | 73 |
| | | GG | 355 | GG | 309 |
| Discovery | case | AA | 0 | TT | 19 |
| | | AG | 44 | GT | 146 |
| | | GG | 233 | GG | 110 |
| Validation | control | AA | 7 | TT | 4 |
| | | AG | 272 | GT | 362 |
| | | GG | 2226 | GG | 2134 |
| Validation | case | AA | 14 | TT | 70 |
| | | AG | 96 | GT | 367 |
| | | GG | 723 | GG | 260 |
| **Total** | | | 4012 | | 3858 |

**Supplementary Table S3. Combined Discovery and Validation Genotypes**, retaining subjects having genotypes of *either* SNP: N=4020.

| | **rs5744168** | | | | **rs35705950** | | | |
|---|---|---|---|---|---|---|---|---|
| | **GG** | **AG** | **AA** | **Total** | **GG** | **GT** | **TT** | **Total** |
| Case | 956 | 140 | 14 | 1110 | 370 | 513 | 89 | 972 |
| Control | 2581 | 304 | 17 | 2902 | 2443 | 435 | 8 | 2886 |
| | Total with rs5744168 data: | | | 4012 | Total with rs35705950 data: | | | 3858 |

**Supplementary Table S4.** Associations with IPF, for rs5744168 (additive and dominant) and rs35705950 (additive) using combined cases from discovery and validation samples, and unscreened non-Finnish European controls from gnomAD.

| **rs5744168** | **cases** | **controls** | **rs35705950** | **cases** | **controls** |
|---|---|---|---|---|---|
| AA | 14 | 258 | TT | 89 | 89 |
| AG | 140 | 7345 | GT | 513 | 1481 |
| GG | 956 | 56941 | GG | 370 | 6135 |
| total | 1110 | 64544 | total | 972 | 7705 |
| Additive OR (95% CI): 1.26 (1.08 – 1.48) p = **0.0041** | | | Additive OR (95% CI): 4.92 (4.37 – 5.54) p < **0.0001** | | |
| Dominant OR (95% CI): 1.21 (1.02 – 1.43) p = **0.0322** | | | | | |

**Supplementary Table S5.** Combined discovery and validation data (subjects having both SNPS: N=3850).

| | rs5744168 | | | rs35705950 | | |
|---|---|---|---|---|---|---|
| | **GG** | **AG** | **AA** | **GG** | **GT** | **TT** |
| Case | 824 | 128 | 12 | 366 | 509 | 89 |
| Control | 2566 | 304 | 16 | 2443 | 435 | 8 |

**Supplementary Table 6**. Logistic regression with interaction of additive rs5744168 and additive rs35705950.

| **Stratification** | **OR (95% CI)** |
|---|---|
| additive rs5744168 in rs35705950 GG stratum | 1.59 (1.21 – 2.07) |
| additive rs5744168 in rs35705950 GT stratum | 1.30 (0.92 – 1.83) |
| additive rs5744168 in rs35705950 TT stratum | 1.07 (0.52 – 2.20) |
| additive rs35705950 in rs5744168 GG stratum | 8.20 (6.93 – 9.71) |
| additive rs35705950 in rs5744168 AG stratum | 6.73 (4.52 – 10.02) |
| additive rs35705950 in rs5744168 AA stratum | 5.52 (2.46 – 12.35) |

**Supplementary Table 7**. Logistic regression with interaction of dominant rs5744168 and additive rs35705950.

| **Stratification** | **OR (95% CI)** |
|---|---|
| dominant rs5744168 in rs35705950 GG stratum | 1.64 (1.22 – 2.21) |
| dominant rs5744168 in rs35705950 GT stratum | 1.25 (0.86 – 1.81) |
| dominant rs5744168 in rs35705950 TT stratum | 0.95 (0.43 – 2.08) |
| additive rs35705950 in rs5744168 GG (non-dominant) stratum | 8.26 (6.98 – 9.79) |
| additive rs35705950 in rs5744168 AG/AA (dominant) stratum | 6.28 (4.08 – 9.68) |

#### Supplementary Table S8. Top 100 enriched gene sets in AEC2s in response to TLR5-agonist.

| NAME | SIZE | ES | NES | NOM p-val | FDR q-val |
|---|---|---|---|---|---|
| GO_RECEPTOR_REGULATOR_ACTIVITY | 325 | 0.18607347 | 3.8065126 | 0 | 0 |
| GO_REGULATION_OF_SIGNALING_RECEPTOR_ACTIVITY | 382 | 0.16077065 | 3.6777642 | 0 | 0 |
| GO_STRUCTURAL_CONSTITUENT_OF_RIBOSOME | 150 | 0.25296393 | 3.561361 | 0 | 0 |
| GO_G_PROTEIN_COUPLED_RECEPTOR_SIGNALING_PATHWAY_COUPLED_TO_CYCLIC_NUCLEOTIDE_SECOND_MESSENGER | 169 | 0.22092628 | 3.4141343 | 0 | 0 |
| GO_CYTOSOLIC_RIBOSOME | 100 | 0.2829385 | 3.335212 | 0 | 0 |
| GO_COTRANSLATIONAL_PROTEIN_TARGETING_TO_MEMBRANE | 92 | 0.28930116 | 3.19563 | 0 | 1.26E-04 |
| GO_EXTRACELLULAR_MATRIX | 409 | 0.13747488 | 3.1821256 | 0 | 1.08E-04 |
| GO_CYTOKINE_ACTIVITY | 131 | 0.23260471 | 3.0869238 | 0 | 2.89E-04 |
| GO_CHEMOKINE_ACTIVITY | 27 | 0.4853821 | 2.9709685 | 0 | 4.25E-04 |
| GO_RIBOSOMAL_SUBUNIT | 177 | 0.19092207 | 2.9340975 | 0 | 6.93E-04 |
| GO_HORMONE_ACTIVITY | 66 | 0.31210375 | 2.9300404 | 0 | 6.30E-04 |
| GO_EXTRACELLULAR_MATRIX_STRUCTURAL_CONSTITUENT | 128 | 0.22112608 | 2.8663318 | 0 | 0.001277872 |
| GO_ESTABLISHMENT_OF_PROTEIN_LOCALIZATION_TO_ENDOPLASMIC_RETICULUM | 104 | 0.23356815 | 2.816181 | 0 | 0.001762545 |
| GO_COLLAGEN_CONTAINING_EXTRACELLULAR_MATRIXGO_COLLAGEN_CONTAINING_EXTRACELLULAR_MATRIX | 325 | 0.13481055 | 2.7814252 | 0 | 0.002452177 |
| GO_NEUROPEPTIDE_RECEPTOR_BINDING | 19 | 0.52601296 | 2.778122 | 0 | 0.002288698 |
| GO_HUMORAL_IMMUNE_RESPONSE | 141 | 0.19965722 | 2.77223 | 0 | 0.002240192 |
| GO_CYTOSOLIC_SMALL_RIBOSOMAL_SUBUNIT | 39 | 0.37512708 | 2.7529147 | 0 | 0.00255696 |
| GO_CHEMOKINE_RECEPTOR_BINDING | 37 | 0.38635007 | 2.7274323 | 0 | 0.00301362 |
| GO_NUCLEAR_TRANSCRIBED_MRNA_CATABOLIC_PROCESS_NONSENSE_MEDIATED_DECAY | 116 | 0.21537103 | 2.710514 | 0 | 0.003416966 |
| GO_RESPONSE_TO_CHEMOKINE | 72 | 0.26468107 | 2.6363358 | 0 | 0.00606049 |
| GO_RESPONSE_TO_BACTERIUM | 449 | 0.10780742 | 2.6328058 | 0 | 0.005879919 |
| GO_EXTERNAL_SIDE_OF_PLASMA_MEMBRANE | 235 | 0.14649744 | 2.6012454 | 0 | 0.007240678 |
| GO_CONDENSED_CHROMOSOME_CENTROMERIC_REGION | 107 | 0.21636058 | 2.5932376 | 0 | 0.007490945 |
| GO_SMALL_RIBOSOMAL_SUBUNIT | 67 | 0.2647012 | 2.570303 | 0 | 0.00848416 |
| GO_MYELOID_LEUKOCYTE_MIGRATION | 158 | 0.16869949 | 2.5328574 | 0 | 0.010824731 |
| GO_ANTIMICROBIAL_HUMORAL_RESPONSE | 59 | 0.28604165 | 2.5255702 | 0 | 0.010849262 |
| GO_REGULATION_OF_SMOOTH_MUSCLE_CONTRACTION | 48 | 0.30396044 | 2.5180566 | 0 | 0.011122458 |
| GO_ANTIMICROBIAL_HUMORAL_IMMUNE_RESPONSE_MEDIATED_BY_ANTIMICROBIAL_PEPTIDE | 34 | 0.3565614 | 2.4803038 | 0 | 0.013906037 |
| GO_G_PROTEIN_COUPLED_RECEPTOR_BINDING | 203 | 0.14756897 | 2.468507 | 0 | 0.014741728 |
| GO_PROTEIN_LOCALIZATION_TO_ENDOPLASMIC_RETICULUM | 128 | 0.18485473 | 2.4657156 | 0 | 0.014478971 |
| GO_SECOND_MESSENGER_MEDIATED_SIGNALING | 347 | 0.11256153 | 2.4478269 | 0 | 0.015415495 |
| GO_CHROMATIN_REMODELING_AT_CENTROMERE | 40 | 0.32579073 | 2.438877 | 0 | 0.016270166 |
| GO_GRANULOCYTE_MIGRATION | 101 | 0.20279466 | 2.4356656 | 0 | 0.016007066 |
| GO_CYTOSOLIC_LARGE_RIBOSOMAL_SUBUNIT | 55 | 0.2734313 | 2.430669 | 0 | 0.016030064 |
| GO_SMOOTH_MUSCLE_CONTRACTION | 84 | 0.22736539 | 2.429463 | 0 | 0.015703212 |
| GO_LARGE_RIBOSOMAL_SUBUNIT | 112 | 0.1957516 | 2.4223733 | 0.001915709 | 0.01609078 |
| GO_CELL_CHEMOTAXIS | 236 | 0.13290162 | 2.3914611 | 0 | 0.0194494 |
| GO_NEUTROPHIL_MIGRATION | 84 | 0.22735123 | 2.3879473 | 0 | 0.019518936 |
| GO_RESPONSE_TO_MOLECULE_OF_BACTERIAL_ORIGIN | 283 | 0.123285785 | 2.3791485 | 0 | 0.020307943 |
| GO_CYCLIC_NUCLEOTIDE_MEDIATED_SIGNALING | 153 | 0.16026226 | 2.370842 | 0 | 0.0211683 |
| GO_AXONEME_ASSEMBLY | 53 | 0.27617428 | 2.3571122 | 0 | 0.022623202 |
| GO_KILLING_OF_CELLS_OF_OTHER_ORGANISM | 22 | 0.41325092 | 2.3501844 | 0 | 0.023209885 |
| GO_CENTROMERE_COMPLEX_ASSEMBLY | 48 | 0.28338844 | 2.3450909 | 0.001976285 | 0.02355524 |
| GO_RESPIRATORY_CHAIN_COMPLEX | 65 | 0.24355203 | 2.3387527 | 0 | 0.024127208 |
| GO_PHOSPHOLIPASE_C_ACTIVATING_G_PROTEIN_COUPLED_RECEPTOR_SIGNALING_PATHWAY | 69 | 0.23855732 | 2.3385031 | 0.001984127 | 0.023625502 |
| GO_EOSINOPHIL_MIGRATION | 15 | 0.49943602 | 2.337546 | 0 | 0.023244595 |
| GO_GLYCOSAMINOGLYCAN_BINDING | 164 | 0.15443222 | 2.3368692 | 0 | 0.022863593 |
| GO_POSITIVE_REGULATION_OF_CELL_ACTIVATION | 271 | 0.12030336 | 2.3259053 | 0.002083333 | 0.02427631 |
| GO_INTERLEUKIN_1_PRODUCTION | 72 | 0.23316878 | 2.3171916 | 0 | 0.02539041 |
| GO_ATTACHMENT_OF_SPINDLE_MICROTUBULES_TO_KINETOCHORE | 29 | 0.36151227 | 2.3069482 | 0 | 0.02646152 |
| GO_CELLULAR_RESPONSE_TO_BIOTIC_STIMULUS | 192 | 0.14300084 | 2.3043265 | 0.00209205 | 0.026479887 |
| GO_MICRO_RIBONUCLEOPROTEIN_COMPLEX | 72 | 0.23098324 | 2.2949321 | 0 | 0.027596468 |
| GO_MRNA_BINDING_INVOLVED_IN_POSTTRANSCRIPTIONAL_GENE_SILENCING | 34 | 0.32984486 | 2.289768 | 0 | 0.02805247 |
| GO_LEUKOCYTE_CHEMOTAXIS | 169 | 0.14440146 | 2.2834108 | 0.00203666 | 0.028844928 |
| GO_REGULATION_OF_VASCULATURE_DEVELOPMENT | 298 | 0.11354177 | 2.2728775 | 0.001953125 | 0.030581892 |
| GO_KINETOCHORE_ORGANIZATION | 20 | 0.4287165 | 2.260823 | 0 | 0.032767855 |
| GO_INTERFERON_GAMMA_PRODUCTION | 91 | 0.20324008 | 2.2573311 | 0.00209205 | 0.032996256 |
| GO_INTERLEUKIN_10_PRODUCTION | 44 | 0.29048693 | 2.2356339 | 0 | 0.037197147 |
| GO_RESPIRASOME | 77 | 0.20994812 | 2.2353542 | 0 | 0.036618195 |
| GO_ATP_SYNTHESIS_COUPLED_ELECTRON_TRANSPORT | 78 | 0.21030737 | 2.214898 | 0.001879699 | 0.041166116 |
| GO_ADENYLATE_CYCLASE_ACTIVATING_G_PROTEIN_COUPLED_RECEPTOR_SIGNALING_PATHWAY | 91 | 0.19774558 | 2.2139015 | 0 | 0.040752146 |
| GO_HUMORAL_IMMUNE_RESPONSE_MEDIATED_BY_CIRCULATING_IMMUNOGLOBULIN | 31 | 0.33159664 | 2.2128477 | 0.003944773 | 0.040413633 |
| GO_ANCHORED_COMPONENT_OF_MEMBRANE | 124 | 0.1682459 | 2.2036152 | 0 | 0.04210628 |
| GO_INTERLEUKIN_1_BETA_PRODUCTION | 61 | 0.23736468 | 2.2028575 | 0 | 0.041685097 |
| GO_MONOCYTE_CHEMOTAXIS | 42 | 0.28509292 | 2.1966262 | 0 | 0.04289445 |
| GO_MATURE_B_CELL_DIFFERENTIATION | 23 | 0.37669465 | 2.187891 | 0.002016129 | 0.044900097 |
| GO_RIBOSOME | 211 | 0.13091859 | 2.1819143 | 0 | 0.04604764 |
| GO_TERTIARY_GRANULE_MEMBRANE | 64 | 0.22335239 | 2.18172 | 0.001937985 | 0.04544833 |
| GO_ADENYLATE_CYCLASE_INHIBITING_G_PROTEIN_COUPLED_RECEPTOR_SIGNALING_PATHWAY | 60 | 0.23746116 | 2.1735845 | 0 | 0.047276147 |
| GO_ACUTE_INFLAMMATORY_RESPONSE | 81 | 0.20298032 | 2.1622891 | 0.003853565 | 0.050477244 |
| GO_SIDE_OF_MEMBRANE | 398 | 0.095704205 | 2.1513705 | 0 | 0.05365028 |
| GO_TERTIARY_GRANULE | 139 | 0.15519238 | 2.1483355 | 0 | 0.05381312 |
| GO_REGULATION_OF_INFLAMMATORY_RESPONSE_TO_ANTIGENIC_STIMULUS | 20 | 0.41378 | 2.1402128 | 0 | 0.05600403 |
| GO_REGULATION_OF_SYNAPTIC_TRANSMISSION_GLUTAMATERGIC | 53 | 0.25328365 | 2.134794 | 0.001960784 | 0.05710955 |
| GO_CONDENSED_CHROMOSOME | 197 | 0.13048762 | 2.132756 | 0.00209205 | 0.05695893 |
| GO_OXIDATIVE_PHOSPHORYLATION | 101 | 0.17958479 | 2.111933 | 0 | 0.064148694 |
| GO_REGULATION_OF_POSTSYNAPTIC_MEMBRANE_POTENTIAL | 102 | 0.18226027 | 2.1064386 | 0 | 0.06558302 |
| GO_REGULATION_OF_HUMORAL_IMMUNE_RESPONSE | 45 | 0.26205274 | 2.0980012 | 0 | 0.068120666 |
| GO_DEFENSE_RESPONSE_TO_BACTERIUM | 138 | 0.1510911 | 2.0929236 | 0 | 0.069359384 |
| GO_CILIUM_MOVEMENT | 53 | 0.2470235 | 2.0922778 | 0.002074689 | 0.06873108 |
| GO_KINETOCHORE | 125 | 0.15830195 | 2.0846431 | 0.00203252 | 0.071476065 |
| GO_CCR_CHEMOKINE_RECEPTOR_BINDING | 19 | 0.39103428 | 2.0801225 | 0.002040816 | 0.072751135 |
| GO_COMPLEMENT_ACTIVATION | 41 | 0.27311864 | 2.0714433 | 0 | 0.07566998 |
| GO_SERINE_HYDROLASE_ACTIVITY | 125 | 0.15950513 | 2.0699213 | 0 | 0.07542111 |
| GO_PROTEIN_LOCALIZATION_TO_KINETOCHORE | 17 | 0.40981698 | 2.051216 | 0 | 0.08351299 |
| GO_SYNAPTIC_TRANSMISSION_GLUTAMATERGIC | 67 | 0.21051715 | 2.050568 | 0.005649718 | 0.08279157 |
| GO_CYTOKINE_RECEPTOR_ACTIVITY | 86 | 0.18924008 | 2.0448759 | 0.00203666 | 0.08494922 |
| GO_CYTOSOLIC_PART | 210 | 0.12058742 | 2.039707 | 0.005940594 | 0.0865996 |
| GO_REGULATION_OF_INFLAMMATORY_RESPONSE | 281 | 0.10536387 | 2.0353572 | 0.006185567 | 0.08768955 |
| GO_DEFENSE_RESPONSE_TO_GRAM_NEGATIVE_BACTERIUM | 39 | 0.28165594 | 2.030215 | 0.003787879 | 0.08938952 |
| GO_DNA_REPLICATION_CHECKPOINT | 15 | 0.4300057 | 2.0259688 | 0.003929273 | 0.09050999 |
| GO_DNA_REPLICATION_INDEPENDENT_NUCLEOSOME_ORGANIZATION | 48 | 0.24160875 | 2.0149872 | 0.003868472 | 0.09551949 |
| GO_U1_SNRNP | 18 | 0.40438315 | 2.0144427 | 0.006465518 | 0.09480532 |
| GO_POSITIVE_REGULATION_OF_LYMPHOCYTE_DIFFERENTIATION | 84 | 0.186088 | 2.0101895 | 0.007722008 | 0.09642225 |
| GO_MEMBRANE_DEPOLARIZATION_DURING_ACTION_POTENTIAL | 31 | 0.2994697 | 2.0026498 | 0.001897533 | 0.099621445 |
| GO_LEUKOCYTE_CELL_CELL_ADHESION | 283 | 0.10222845 | 2.001681 | 0.00210084 | 0.09913249 |
| GO_G_PROTEIN_COUPLED_RECEPTOR_ACTIVITY | 264 | 0.10809335 | 1.9997804 | 0.001945525 | 0.09930617 |
| GO_REGULATION_OF_SYSTEM_PROCESS | 458 | 0.08162216 | 1.9995226 | 0.003913894 | 0.09846414 |
| GO_ACUTE_PHASE_RESPONSE | 31 | 0.3039531 | 1.9960674 | 0.007858546 | 0.09949394 |
| GO_THREONINE_TYPE_PEPTIDASE_ACTIVITY | 20 | 0.36935124 | 1.9882864 | 0.006329114 | 0.1031309 |

### Supplementary Table S9. Top 100 enriched gene sets in bronchial epithelial cells in response to TLR5-agonist.

| NAME | SIZE | ES | NES | NOM p-val | FDR q-val |
|---|---|---|---|---|---|
| GO_CILIUM | 611 | 0.26462924 | 7.4490876 | 0 | 0 |
| GO_CILIUM_ORGANIZATION | 384 | 0.30512163 | 6.9362035 | 0 | 0 |
| GO_AXONEME_ASSEMBLY | 68 | 0.6333772 | 6.1527853 | 0 | 0 |
| GO_CELL_PROJECTION_ASSEMBLY | 567 | 0.22212964 | 6.132419 | 0 | 0 |
| GO_EXTRACELLULAR_MATRIX | 512 | 0.2278935 | 5.9158325 | 0 | 0 |
| GO_CILIUM_MOVEMENT | 138 | 0.42300472 | 5.8055744 | 0 | 0 |
| GO_CELL_JUNCTION_ORGANIZATION | 680 | 0.19349155 | 5.714895 | 0 | 0 |
| GO_CILIARY_PLASM | 121 | 0.4377222 | 5.605424 | 0 | 0 |
| GO_UROGENITAL_SYSTEM_DEVELOPMENT | 312 | 0.27449447 | 5.4931817 | 0 | 0 |
| GO_MICROTUBULE_BUNDLE_FORMATION | 99 | 0.46570683 | 5.4476604 | 0 | 0 |
| GO_SENSORY_ORGAN_DEVELOPMENT | 522 | 0.20556803 | 5.407369 | 0 | 0 |
| GO_CELL_CELL_JUNCTION | 471 | 0.21211599 | 5.316641 | 0 | 0 |
| GO_MOTILE_CILIUM | 178 | 0.3225095 | 5.122909 | 0 | 0 |
| GO_COLLAGEN_CONTAINING_EXTRACELLULAR_MATRIX | 391 | 0.20961045 | 4.8652616 | 0 | 0 |
| GO_CELL_CELL_ADHESION_VIA_PLASMA_MEMBRANE_ADHESION_MOLECULES | 241 | 0.26595068 | 4.852057 | 0 | 0 |
| GO_REGULATION_OF_MEMBRANE_POTENTIAL | 406 | 0.21249594 | 4.851823 | 0 | 0 |
| GO_CALCIUM_ION_BINDING | 630 | 0.17180537 | 4.847991 | 0 | 0 |
| GO_CELL_JUNCTION_ASSEMBLY | 414 | 0.20626739 | 4.83569 | 0 | 0 |
| GO_KIDNEY_EPITHELIUM_DEVELOPMENT | 132 | 0.3524084 | 4.8128185 | 0 | 0 |
| GO_MICROTUBULE_BASED_MOVEMENT | 344 | 0.22276054 | 4.7463617 | 0 | 0 |
| GO_EXTRACELLULAR_MATRIX_STRUCTURAL_CONSTITUENT | 150 | 0.32630318 | 4.691233 | 0 | 0 |
| GO_GATED_CHANNEL_ACTIVITY | 301 | 0.23408452 | 4.689806 | 0 | 0 |
| GO_PATTERN_SPECIFICATION_PROCESS | 397 | 0.20285106 | 4.656498 | 0 | 0 |
| GO_AXONEMAL_DYNEIN_COMPLEX_ASSEMBLY | 33 | 0.6865023 | 4.6124053 | 0 | 0 |
| GO_HOMOPHILIC_CELL_ADHESION_VIA_PLASMA_MEMBRANE_ADHESION_MOLECULES | 139 | 0.33702016 | 4.5579 | 0 | 0 |
| GO_SENSORY_ORGAN_MORPHOGENESIS | 247 | 0.2512412 | 4.5528607 | 0 | 0 |
| GO_MESONEPHROS_DEVELOPMENT | 95 | 0.4016111 | 4.551198 | 0 | 0 |
| GO_CELL_MORPHOGENESIS_INVOLVED_IN_NEURON_DIFFERENTIATION | 573 | 0.16417737 | 4.5394816 | 0 | 0 |
| GO_SYNAPSE_ORGANIZATION | 406 | 0.19451457 | 4.5210257 | 0 | 0 |
| GO_RESPIRATORY_SYSTEM_DEVELOPMENT | 198 | 0.2742395 | 4.515719 | 0 | 0 |
| GO_CILIARY_BASAL_BODY | 151 | 0.31915528 | 4.495792 | 0 | 0 |
| GO_TISSUE_MORPHOGENESIS | 629 | 0.16037902 | 4.482496 | 0 | 0 |
| GO_SENSORY_SYSTEM_DEVELOPMENT | 357 | 0.20336844 | 4.462335 | 0 | 0 |
| GO_EMBRYONIC_ORGAN_MORPHOGENESIS | 277 | 0.23073773 | 4.4481473 | 0 | 0 |
| GO_APICAL_JUNCTION_COMPLEX | 137 | 0.32043925 | 4.432336 | 0 | 0 |
| GO_CELLULAR_COMPONENT_MORPHOGENESIS | 757 | 0.14128038 | 4.4116573 | 0 | 0 |
| GO_CELL_PART_MORPHOGENESIS | 669 | 0.14902945 | 4.4088173 | 0 | 0 |
| GO_EMBRYONIC_MORPHOGENESIS | 555 | 0.16918966 | 4.406399 | 0 | 0 |
| GO_EXTRACELLULAR_STRUCTURE_ORGANIZATION | 368 | 0.19950876 | 4.3641887 | 0 | 0 |
| GO_POSTSYNAPTIC_SPECIALIZATION_MEMBRANE | 108 | 0.35632727 | 4.361286 | 0 | 0 |
| GO_BRANCHING_MORPHOGENESIS_OF_AN_EPITHELIAL_TUBE | 144 | 0.31092972 | 4.357689 | 0 | 0 |
| GO_SYNAPSE_ASSEMBLY | 170 | 0.2865472 | 4.35526 | 0 | 0 |
| GO_MORPHOGENESIS_OF_A_BRANCHING_STRUCTURE | 187 | 0.27884695 | 4.344144 | 0 | 0 |
| GO_CELL_MORPHOGENESIS_INVOLVED_IN_DIFFERENTIATION | 713 | 0.14320879 | 4.3277445 | 0 | 0 |
| GO_MORPHOGENESIS_OF_AN_EPITHELIUM | 529 | 0.16386792 | 4.3255897 | 0 | 0 |
| GO_EPITHELIAL_CELL_DIFFERENTIATION | 612 | 0.152715 | 4.30264 | 0 | 0 |
| GO_CILIUM_OR_FLAGELLUM_DEPENDENT_CELL_MOTILITY | 104 | 0.36022723 | 4.2861605 | 0 | 0 |
| GO_CAMERA_TYPE_EYE_DEVELOPMENT | 302 | 0.21178058 | 4.25798 | 0 | 0 |
| GO_CATION_CHANNEL_COMPLEX | 206 | 0.25024796 | 4.2111154 | 0 | 0 |
| GO_NEURON_TO_NEURON_SYNAPSE | 354 | 0.19379352 | 4.1731086 | 0 | 0 |
| GO_TRANSPORTER_COMPLEX | 303 | 0.20985423 | 4.159187 | 0 | 0 |
| GO_EPITHELIAL_TUBE_MORPHOGENESIS | 314 | 0.20072545 | 4.1516457 | 0 | 0 |
| GO_EXTRACELLULAR_MATRIX_BINDING | 55 | 0.46172115 | 4.131769 | 0 | 0 |
| GO_AXONEMAL_DYNEIN_COMPLEX | 19 | 0.8040414 | 4.123685 | 0 | 0 |
| GO_POSTSYNAPTIC_MEMBRANE | 255 | 0.2224701 | 4.097058 | 0 | 0 |
| GO_APICAL_PLASMA_MEMBRANE | 335 | 0.19637586 | 4.0795116 | 0 | 0 |
| GO_CIRCULATORY_SYSTEM_PROCESS | 513 | 0.15713742 | 4.072833 | 0 | 0 |
| GO_APICAL_PART_OF_CELL | 399 | 0.17633456 | 4.061084 | 0 | 0 |
| GO_SYNAPTIC_SIGNALING | 681 | 0.14013985 | 4.0512323 | 0 | 0 |
| GO_PASSIVE_TRANSMEMBRANE_TRANSPORTER_ACTIVITY | 425 | 0.17328629 | 4.0233307 | 0 | 0 |
| GO_TIGHT_JUNCTION | 123 | 0.3083854 | 4.0174713 | 0 | 0 |
| GO_PLASMA_MEMBRANE_BOUNDED_CELL_PROJECTION_CYTOPLASM | 207 | 0.23920767 | 4.0088253 | 0 | 0 |
| GO_REGIONALIZATION | 308 | 0.19590433 | 4.0006204 | 0 | 0 |
| GO_REGULATION_OF_SYSTEM_PROCESS | 550 | 0.14790955 | 3.997414 | 0 | 0 |
| GO_EAR_DEVELOPMENT | 204 | 0.23673886 | 3.9920428 | 0 | 0 |
| GO_SENSORY_PERCEPTION | 510 | 0.15294094 | 3.9553277 | 0 | 0 |
| GO_EMBRYONIC_ORGAN_DEVELOPMENT | 418 | 0.16933458 | 3.9442036 | 0 | 0 |
| GO_TRANSMEMBRANE_RECEPTOR_PROTEIN_TYROSINE_KINASE_ACTIVITY | 60 | 0.43322346 | 3.9295857 | 0 | 0 |
| GO_REGULATION_OF_HORMONE_LEVELS | 457 | 0.15646644 | 3.9086354 | 0 | 0 |
| GO_POSTSYNAPTIC_DENSITY_MEMBRANE | 84 | 0.35979292 | 3.9078102 | 0 | 0 |
| GO_APPENDAGE_DEVELOPMENT | 162 | 0.2622846 | 3.8937159 | 0 | 0 |
| GO_APPENDAGE_MORPHOGENESIS | 133 | 0.29195145 | 3.891496 | 0 | 0 |
| GO_KIDNEY_MORPHOGENESIS | 87 | 0.35300344 | 3.866379 | 0 | 0 |
| GO_SYNAPTIC_MEMBRANE | 360 | 0.17731404 | 3.8592052 | 0 | 0 |
| GO_EPIDERMIS_DEVELOPMENT | 329 | 0.18520296 | 3.85728 | 0 | 0 |
| GO_ACTIN_FILAMENT_BASED_PROCESS | 764 | 0.12263943 | 3.8563368 | 0 | 0 |
| GO_ADHERENS_JUNCTION | 165 | 0.2564715 | 3.8141146 | 0 | 0 |
| GO_NEPHRON_EPITHELIUM_DEVELOPMENT | 100 | 0.32495102 | 3.81391 | 0 | 0 |
| GO_PROTEIN_TRANSPORT_ALONG_MICROTUBULE | 68 | 0.38949025 | 3.8132586 | 0 | 0 |
| GO_EYE_MORPHOGENESIS | 145 | 0.2662824 | 3.8112056 | 0 | 0 |
| GO_VOLTAGE_GATED_ION_CHANNEL_ACTIVITY | 180 | 0.24291204 | 3.790168 | 0 | 0 |
| GO_NEPHRON_DEVELOPMENT | 133 | 0.28429705 | 3.7849908 | 0 | 0 |
| GO_CELL_CELL_JUNCTION_ORGANIZATION | 203 | 0.23050746 | 3.7676747 | 0 | 0 |
| GO_PLASMA_MEMBRANE_PROTEIN_COMPLEX | 527 | 0.14405295 | 3.7675335 | 0 | 0 |
| GO_CYTOPLASMIC_REGION | 249 | 0.20596449 | 3.7485216 | 0 | 0 |
| GO_METAL_ION_TRANSMEMBRANE_TRANSPORTER_ACTIVITY | 398 | 0.16310132 | 3.7484534 | 0 | 0 |
| GO_TRANSMEMBRANE_RECEPTOR_PROTEIN_KINASE_ACTIVITY | 78 | 0.36413926 | 3.7457595 | 0 | 0 |
| GO_EMBRYONIC_APPENDAGE_MORPHOGENESIS | 112 | 0.30387774 | 3.7427444 | 0 | 0 |
| GO_RECEPTOR_REGULATOR_ACTIVITY | 414 | 0.16251594 | 3.7359588 | 0 | 0 |
| GO_INTRACILIARY_TRANSPORT | 53 | 0.43395075 | 3.7292871 | 0 | 0 |
| GO_RENAL_TUBULE_DEVELOPMENT | 87 | 0.341448 | 3.7073388 | 0 | 0 |
| GO_SKIN_DEVELOPMENT | 282 | 0.19352163 | 3.6942763 | 0 | 0 |
| GO_9PLUS2_MOTILE_CILIUM | 104 | 0.29866567 | 3.6868632 | 0 | 0 |
| GO_REGULATION_OF_BLOOD_CIRCULATION | 271 | 0.19444543 | 3.6754527 | 0 | 0 |
| GO_DIGESTIVE_SYSTEM_DEVELOPMENT | 129 | 0.27983227 | 3.6621695 | 0 | 0 |
| GO_INTRACILIARY_TRANSPORT_PARTICLE | 27 | 0.5981022 | 3.6466236 | 0 | 0 |
| GO_CILIARY_TRANSITION_ZONE | 67 | 0.37858054 | 3.638657 | 0 | 0 |
| GO_POSITIVE_REGULATION_OF_SYNAPSE_ASSEMBLY | 61 | 0.40267494 | 3.6247911 | 0 | 0 |
| GO_CATION_CHANNEL_ACTIVITY | 301 | 0.18187964 | 3.624447 | 0 | 0 |
| GO_REGULATION_OF_TRANS_SYNAPTIC_SIGNALING | 426 | 0.15498443 | 3.62422 | 0 | 0 |

**Supplementary Table S10.** ***Tlr5*-deficient lung epithelia fail to upregulate antimicrobial factors in response to TLR5-agonism.**

| | *p-value* | |
|---|---|---|
| **Name** | ***Tlr5Fx/Fx*** | ***Tlr5Fx/Fx SPC-Cre*** |
| c001.Immune_Response-GO_ANTIMICROBIAL_HUMORAL_IMMUNE_RESPONSE_MEDIATED_BY_ANTIMICROBIAL_PEPTIDE | **0.008924108** | 0.267169234 |
| c001.Immune_Response-GO_ANTIMICROBIAL_HUMORAL_RESPONSE | **0.008321884** | 0.372860842 |
| c001.Immune_Response-GO_HUMORAL_IMMUNE_RESPONSE | **0.003036545** | 0.098463875 |
| c002.Leukocyte_Migration-GO_EOSINOPHIL_MIGRATION | **0.019034289** | 0.100907044 |
| c002.Leukocyte_Migration-GO_GRANULOCYTE_MIGRATION | **0.022583681** | 0.088046684 |
| c002.Leukocyte_Migration-GO_MYELOID_LEUKOCYTE_MIGRATION | **0.024379226** | 0.125033277 |
| c002.Leukocyte_Migration-GO_NEUTROPHIL_MIGRATION | **0.023989337** | 0.083508658 |
| c003.Response_to_Stimulus-GO_CELL_CHEMOTAXIS | **0.034108574** | 0.179770223 |
| c003.Response_to_Stimulus-GO_CHEMOKINE_ACTIVITY | **0.004606979** | **0.031377117** |
| c003.Response_to_Stimulus-GO_CYCLIC_NUCLEOTIDE_MEDIATED_SIGNALING | **0.014586506** | 0.20202499 |
| c003.Response_to_Stimulus-GO_CYTOKINE_ACTIVITY | **0.013516942** | 0.097277202 |
| c003.Response_to_Stimulus-GO_RESPONSE_TO_BACTERIUM | **0.002902054** | 0.046970232 |
| c003.Response_to_Stimulus-GO_RESPONSE_TO_CHEMOKINE | **0.0089151** | 0.132150558 |
| c003.Response_to_Stimulus-GO_RESPONSE_TO_MOLECULE_OF_BACTERIAL_ORIGIN | **0.005380296** | 0.068274296 |
| c004.Extracellular_Matrix-GO_EXTRACELLULAR_MATRIX_STRUCTURAL_CONSTITUENT | **0.00502209** | 0.013259411 |
| c004.Extracellular_Matrix-GO_EXTRACELLULAR_MATRIX | **0.003524195** | 0.040737876 |
| c005.Cell_Killing-GO_KILLING_OF_CELLS_OF_OTHER_ORGANISM | **0.028849519** | 0.543053869 |
| c006.Plasma_membrane_bounded_cell_projection_organization-GO_AXONEME_ASSEMBLY | 0.076764719 | 0.28826602 |
| c007.Chemokine_receptor_binding-GO_CHEMOKINE_RECEPTOR_BINDING | **0.005268301** | **0.046097586** |
| c008.Cytokine_production-GO_INTERLEUKIN_1_PRODUCTION | **0.032549379** | 0.329205066 |
| c009.Carbohydrate_derivative_binding-GO_GLYCOSAMINOGLYCAN_BINDING | **0.002190312** | **0.016635391** |
| c010.Signaling_transduction-GO_G_PROTEIN_COUPLED_RECEPTOR_BINDING | **0.014772434** | 0.076623897 |
| c010.Signaling_transduction-GO_G_PROTEIN_COUPLED_RECEPTOR_SIGNALING_PATHWAY_COUPLED_TO_CYCLIC_NUCLEOTIDE_SECOND_MESSENGER | **0.0137268** | 0.204618423 |
| c010.Signaling_transduction-GO_PHOSPHOLIPASE_C_ACTIVATING_G_PROTEIN_COUPLED_RECEPTOR_SIGNALING_PATHWAY | **0.030680511** | 0.363229477 |
| c010.Signaling_transduction-GO_RECEPTOR_REGULATOR_ACTIVITY | **0.017040135** | 0.064047113 |
| c010.Signaling_transduction-GO_REGULATION_OF_SIGNALING_RECEPTOR_ACTIVITY | **0.018453848** | 0.091614756 |
| c010.Signaling_transduction-GO_SECOND_MESSENGER_MEDIATED_SIGNALING | **0.021113035** | 0.21154008 |
| c011.Ribosome-GO_CYTOSOLIC_LARGE_RIBOSOMAL_SUBUNIT | 0.244530362 | 0.308305924 |
| c011.Ribosome-GO_CYTOSOLIC_RIBOSOME | 0.24635125 | 0.298188006 |
| c011.Ribosome-GO_CYTOSOLIC_SMALL_RIBOSOMAL_SUBUNIT | 0.251133522 | 0.291329955 |
| c011.Ribosome-GO_LARGE_RIBOSOMAL_SUBUNIT | 0.217212844 | 0.307197216 |
| c011.Ribosome-GO_RIBOSOMAL_SUBUNIT | 0.234311734 | 0.297388483 |
| c011.Ribosome-GO_SMALL_RIBOSOMAL_SUBUNIT | 0.242297818 | 0.284749853 |
| c011.Ribosome-GO_STRUCTURAL_CONSTITUENT_OF_RIBOSOME | 0.2485434 | 0.300530047 |
| c012.Chromosome-GO_ATTACHMENT_OF_SPINDLE_MICROTUBULES_TO_KINETOCHORE | 0.224036999 | 0.092568829 |
| c012.Chromosome-GO_CENTROMERE_COMPLEX_ASSEMBLY | 0.104832053 | 0.110484605 |
| c012.Chromosome-GO_CHROMATIN_REMODELING_AT_CENTROMERE | 0.067413456 | 0.14563414 |
| c012.Chromosome-GO_CONDENSED_CHROMOSOME_CENTROMERIC_REGION | 0.177360936 | 0.130360252 |
| c013.Respirasome-GO_RESPIRATORY_CHAIN_COMPLEX | 0.168437319 | 0.218494102 |
| c014.Other-GO_COTRANSLATIONAL_PROTEIN_TARGETING_TO_MEMBRANE | 0.230460958 | 0.301770036 |
| c014.Other-GO_ESTABLISHMENT_OF_PROTEIN_LOCALIZATION_TO_ENDOPLASMIC_RETICULUM | 0.227083172 | 0.312011933 |
| c014.Other-GO_EXTERNAL_SIDE_OF_PLASMA_MEMBRANE | **0.033757611** | 0.289190889 |
| c014.Other-GO_HORMONE_ACTIVITY | **0.017760449** | 0.11026162 |
| c014.Other-GO_NEUROPEPTIDE_RECEPTOR_BINDING | 0.175883279 | 0.215185567 |
| c014.Other-GO_NUCLEAR_TRANSCRIBED_MRNA_CATABOLIC_PROCESS_NONSENSE_MEDIATED_DECAY | 0.212817069 | 0.299901195 |
| c014.Other-GO_POSITIVE_REGULATION_OF_CELL_ACTIVATION | **0.022066384** | 0.203019109 |
| c014.Other-GO_PROTEIN_LOCALIZATION_TO_ENDOPLASMIC_RETICULUM | 0.209424858 | 0.316895001 |
| c014.Other-GO_REGULATION_OF_SMOOTH_MUSCLE_CONTRACTION | **0.012816058** | 0.141872677 |
| c014.Other-GO_SMOOTH_MUSCLE_CONTRACTION | **0.011279759** | 0.114403396 |
